# Two CTCF motifs impede cohesin-mediated DNA loop extrusion

**DOI:** 10.1101/2025.01.26.634934

**Authors:** Roman Barth, Richard Janissen, Laura Muras, Jaco van der Torre, Gabriele Litos, Eli van der Sluis, Ashmiani van der Graaf, Iain F. Davidson, Jan-Michael Peters, Cees Dekker

**Author notes:** These authors contributed equally to this work (Roman Barth, Richard Janissen, Laura Muras). Institute of Protein Design, Department of Biochemistry, Seattle, USA. Institute of Bioengineering, Deggendorf Institute of Technology; Oberschneiding, 94363, Germany (Richard Janissen). Center for Plant Molecular Biology (ZMBP), University of Tübingen, Tübingen, 72076, Germany.

## Abstract

Cohesin extrudes DNA into loops and is positioned along the genome by stalling at CTCF upon encountering its N-terminal region (NTR). The mechanism underlying this stalling, however, is unresolved. Using single-molecule assays that monitor DNA loop extrusion (LE) in the presence of NTR fragments, we identify two amino acid motifs, YDF and KTYQR, that hinder LE. KTYQR is found to fully impede LE activity, while YDF hinders cohesin to complete LE step cycles and converts cohesin into a unidirectional extruder by strengthening the affinity of STAG1 to DNA. We thus identify two distinct NTR motifs that stall LE via different yet synergistic mechanisms, highlighting the multifaceted ways employed by CTCF to modulate LE to shape and regulate genomes.

**One-Sentence Summary:** The N-terminus of CTCF employs two independent motifs that synergistically stall cohesin-mediated DNA loop extrusion.

---

The spatial organization of the genome is crucial for processes like transcription, replication, and DNA repair (*1*, *2*). Central to the chromosomal architecture are DNA loops that are progressively extruded by the Structural Maintenance of Chromosomes (SMC) ATPases condensin I/II, cohesin, and SMC5/6 in humans (*3*–*10*). Cohesin (Fig. 1A) forms DNA loops in interphase (*11*–*14*) and extrudes DNA asymmetrically while frequently switching direction (*3*, *15*), forming large topologically associated domains (TADs) whose boundaries are demarcated by CCCTC-binding factor (CTCF) (*12*, *14*, *16*–*20*). CTCF positions DNA loops at TAD boundaries by stalling cohesin, a tightly regulated process that is directional: The majority of CTCF binding sites are oriented such that CTCF’s N-terminal region (NTR) points toward the TAD interior (*14*, *21*–*25*), indicating that the NTR mediates loop positioning. Indeed, previous single-molecule *in vitro* experiments showed that CTCF stalls DNA loop extrusion (LE) by cohesin particularly when approached from its NTR (*26*, *27*). The interaction depends strongly, but not exclusively, on a YDF motif (*26*, *28*–*32*) in the NTR (residues 226-228 of CTCF; Fig. 1A) that binds to a ‘conserved essential surface’ (CES) formed by the kleisin and STAG1/2 cohesin subunits (*33*–*39*). Mutations in the YDF motif reduce CTCF-anchored loops (*33*) but have less impact on cohesin enrichment at CTCF sites (*33*), indicating that cohesin-CTCF co-localization alone is insufficient to form these loops (*29*, *33*, *40*–*42*) or that the YDF motif isn’t the only mechanism by which cohesin stalls at CTCF sites. The underlying mechanism of how CTCF inhibits cohesin-mediated LE has been coined a ‘mechanistic mystery’ (*40*). Indeed, it remains an important unresolved question of how CTCF regulates cohesin-driven LE and hence impacts critical cellular processes such as protocadherin isoform diversity (*43*), diseases like cohesinopathies or cancer (*44*), and in V(D)J recombination (*45*, *46*).

**Figure 1.**
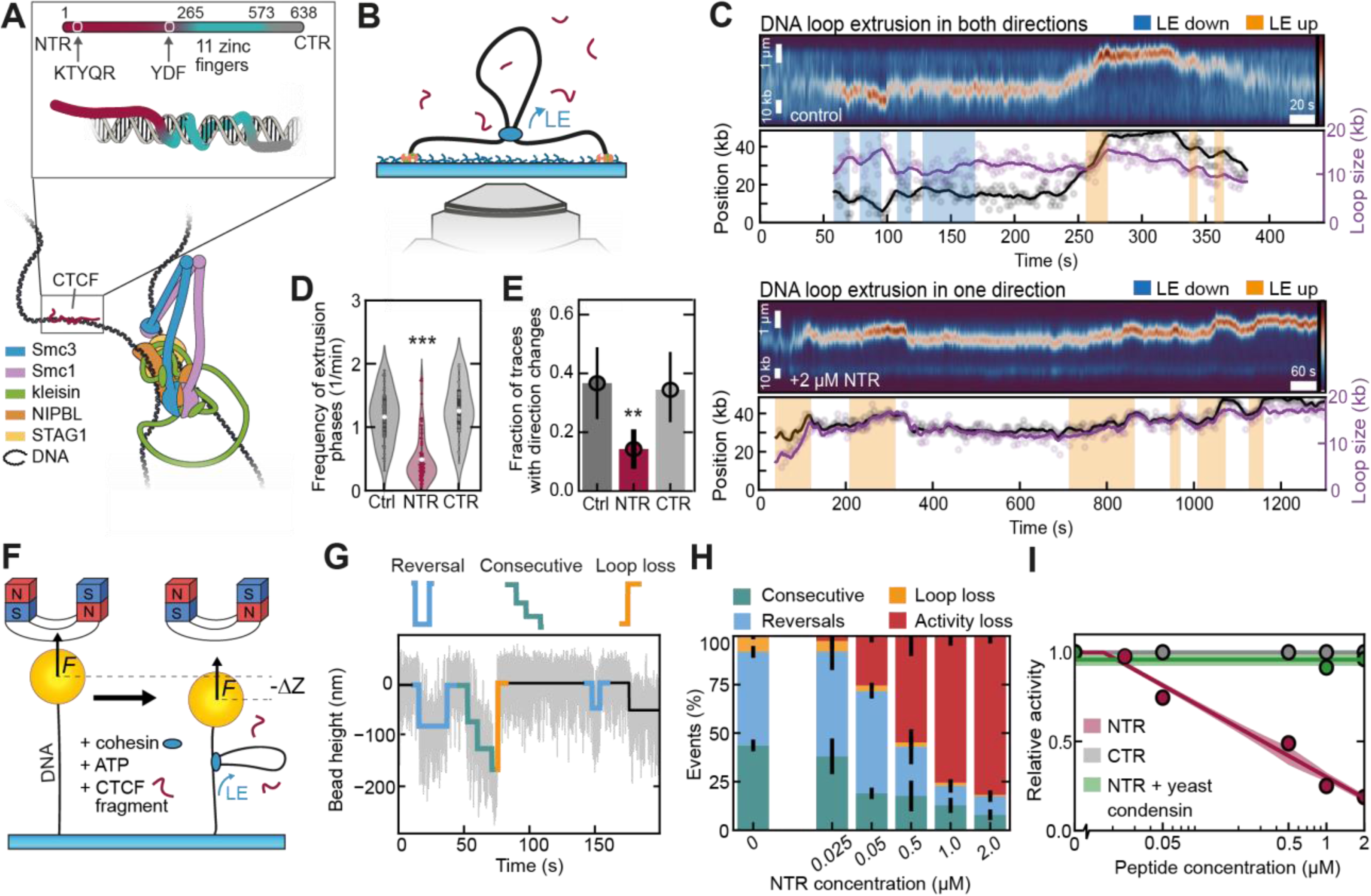
The N-terminal region of CTCF impedes cohesin-mediated loop extrusion. (**A**) Illustration of a DNA-bound human cohesin complex (bottom; based on PDB 7W1M (*27*)) and CTCF (top). CTCF shows the N-terminal region (NTR; red) and C-terminal region (CTR; grey) of CTCF, separated by 11 zinc fingers (cyan) that bind to DNA. (**B**) Fluorescence visualisation assay to observe cohesin-mediated loop extrusion on individual, surface-tethered DNA molecules. (**C**) Example kymograph of LE that consistently extrudes in both directions (lower panel) or towards one direction (upper panel). Segmented extrusion phases (Methods and (*15*)) are shaded yellow for extrusion upwards and blue for extrusion downwards. Loop position (black) and size (purple) are shown below. Solid lines depict smoothed data (Methods). (**D**) Violin plot of the frequency with which extrusion phases occur in the absence (control) or the presence of 2 μM CTCF NTR or CTR. White dots denote the mean, the box shows the quartiles of the data, and whiskers extend to 1.5 * interquartile range. Statistical significance was assessed by a Kruskal-Wallis test (***: p < 0.001 and the Cliff’s Delta statistic is ≥ 0.33). N = 45, 91, 40 DNA molecules from left to right. (**E**) Fraction of traces that show direction changes in the absence (control) or the presence of 2 μM CTCF NTR or CTR. Error bar denotes the 95% binomial confidence interval. Statistical significance was assessed by a two-sided Fisher’s exact test (**: p < 0.01 and the 95% odds ratio CI does not contain 1). N = 58, 103, 57 DNA molecules from left to right. (**F**) Magnetic Tweezers (MT) experimental setup used to measure individual LE steps (*26*, *47*, *49*). (**G**) Example MT trace and illustration of the classification of LE stepping events into reversals (blue), consecutive LE steps (green), and loop loss (orange). Raw data is shown in light grey, the black line shows the resulting fit from the stepfinding algorithm (Methods). (**H**) Fraction of events classified as consecutive steps (green), reversals (blue), and loop loss (orange) for MT traces with increasing concentrations of CTCF NTR. Loss in activity (red) denotes that proportionally fewer steps are observed compared to the control measurement ([NTR] = 0 μM). Error bars denote 95% binomial confidence intervals (CI). N = 1097, 109, 537, 46, 75, 77 steps from 193, 19, 114, 16, 51, 103 tethers, respectively, for increasing CTCF NTR concentration. (**I**) The relative stepping activity of cohesin, compared to the corresponding control ([peptide] = 0 μM), for increasing concentrations of CTCF NTR, CTR, and NTR in combination with yeast condensin. The data corresponds to the inverse of the red bar height of panels H and Fig. S1K,L, respectively. Fits serve as a guide for the eye (Methods). Lines denote the fit and the shaded error bars denote the mean and interpolated SD of the individual data points.

With the aim to understand the molecular mechanisms by which CTCF affects cohesin-mediated LE, we dissected the CTCF NTR into fragments and investigated their effects on LE with single-molecule and bulk biochemical methods. We found that the presence of the complete NTR reduces the frequency of active LE phases and caused cohesin to mainly extrude loops from one side only. The latter effect is attributed to an increase of STAG1’s DNA affinity induced by the YDF motif, preventing DNA strand exchange and direction switching. While the YDF motif reduces cohesin’s ATPase activity as well as hinders cohesin in taking consecutive LE steps and, its effect on cohesin is weaker than the full NTR. We identified and studied a second motif, KTYQR (residues 23-27 of CTCF (*41*)), that independently reduces cohesin’s ATPase activity and blocks cohesin-mediated LE. These findings clarify how CTCF influences cohesin’s LE dynamics and shapes genomes.

## The NTR of CTCF impedes cohesin-mediated DNA loop extrusion

To probe cohesin’s LE behavior in response to the CTCF NTR and C-terminal region (CTR) (Fig. S1A) on the single-molecule level, we added purified 100 pM human cohesin^STAG1^ (henceforth ‘cohesin’) and 250 pM NIPBL-Mau2 to surface-tethered λ-DNA (*4*) (Fig. 1B; Methods), and measured its LE activity in the presence of a buffer containing fragments of CTCF. This approach probes loop extrusion by cohesin in a buffer with CTCF fragments in solution, which provides a high-throughput assay that probes the essential interactions between cohesin and CTCF which in cells are both bound on DNA. Since short amino acid sequences often bind to their interactions partners with lower affinity as isolated peptides than in the context of folded proteins, we used CTCF peptides in large excess over cohesin and controlled their binding specificities by using scrambled and mutated peptides. The LE dynamics in kymographs (Fig. 1C; Methods) show that, as reported before (*15*), cohesin alternated between active extrusion phases and diffusion/slipping phases, with stochastic direction changes (Fig. 1C) occurring upon exchange of the NIPBL subunit. In the presence of 2 μM NTR (residues 2-259) in this single-molecule visualization assay, the frequency of active extrusion phases decreased ∼3-fold compared to conditions without NTR (Fig. 1D). Notably, the extrusion rate during active phases remained unchanged (Fig. S1B), suggesting that the NTR particularly hinders cohesin to initiate LE phases. Bidirectional extrusion (i.e. that cohesin sequentially reels in DNA from left and from right side during traces) decreased from 35% to 12% of cases in the presence of NTR (Fig. 1E), indicating that the NTR converts cohesin into a predominantly unidirectional extruder (reeling in DNA consequently from one side only). The CTR had no effect on the LE activity or directionality.

Magnetic tweezers (MT) experiments (Fig. 1F; Methods), which are able to detect LE on individual DNA molecules with single LE step resolution (*26*, *47*–*49*), showed three distinct signatures (Fig. 1G): consecutive downward steps (>1 step), single downward steps followed by upward steps of the same size (‘reversals’; Methods; (*47*, *49*)), and (rare) large upward steps that resolved previous downward steps (‘loop loss’). Consecutive steps relate to cohesin’s stepwise LE activity (*26*, *47*, *49*) while we attribute reversals to incomplete LE step cycles, e.g. as a consequence of the inability to hydrolyze ATP (*47*, *49*). Current mechanistic models propose that DNA is reeled into the Smc lumen upon ATP binding, creating a downward step (*50*–*52*). Recent molecular dynamics simulations of SMC-driven LE suggest that DNA may often slip out of the lumen (*53*), constituting unsuccessful LE steps. We therefore consider reversals as failed attempts at completing the LE hydrolysis cycle (Fig. S1C). We developed a step-finding algorithm which more accurately extracted steps compared to our previous method (Methods; Fig. S1D-H (*54*)).

In presence of the NTR of CTCF, cohesin’s LE activity decreased in a concentration-dependent manner, with fewer steps observed within our 12-minute acquisition period (Fig. 1H; Fig. S1I,J), where the total activity dropped to 20% at 2 μM NTR compared to control experiments without NTR. Both consecutive steps and reversals were equally affected, maintaining a constant ratio (Fig. S1M-O). Importantly, no dependence on CTCF was observed in similar experiments using yeast condensin (Fig. S1K,M-O; Fig. 1I; Methods), confirming the specificity of NTR’s impact on human cohesin. Similarly, addition of the CTR (Fig. 1I and Fig. S1L-O) had no effect whatsoever, aligning with recent studies that found that the NTR, not the CTR interacts with cohesin (*26*, *27*, *31*, *33*, *41*, *55*). We conclude that the CTCF NTR biases cohesin toward unidirectional extrusion (Fig. 1E) and significantly reduces cohesin’s stepping activity in a concentration-dependent manner (Fig. 1D,I).

## The YDF motif biases cohesin towards unidirectional extrusion

The CTCF NTR contains a conserved YDF motif (residues 226-228; Fig. 2A; Fig. S2A) that binds the CES on cohesin’s STAG1/2 subunit (*33*–*39*) that was observed to impact TAD formation (*33*). To study the role of the YDF motif in inhibiting LE by cohesin, we used a very short NTR fragment containing the YDF motif (residues 222-232; ‘YDF peptide’) and a mutated version (‘ADA peptide’; Fig. 2B; Table S1). Increasing concentrations of the YDF peptide significantly reduced cohesin’s ATPase activity (Fig. 2C; Methods) and decreased the frequency of active extrusion phases (Fig. 2D) by 1.5-fold, whereas the ADA peptide had no effect on either activity. Variants of the full CTCF NTR with YDF mutations (NTR^FDF^ and NTR^ADA^; Fig. 2B; Fig. S3A; Table S1) reduced the frequency of extrusion phases by 2-fold (Fig. 2D) but did not affect the extrusion rate (Fig. S3B). The partial inhibition of cohesin observed with the NTR^ADA^ fragment, but not the ADA peptide, suggests that additional motifs, beyond YDF, contribute to cohesin regulation by CTCF.

**Figure 2.**
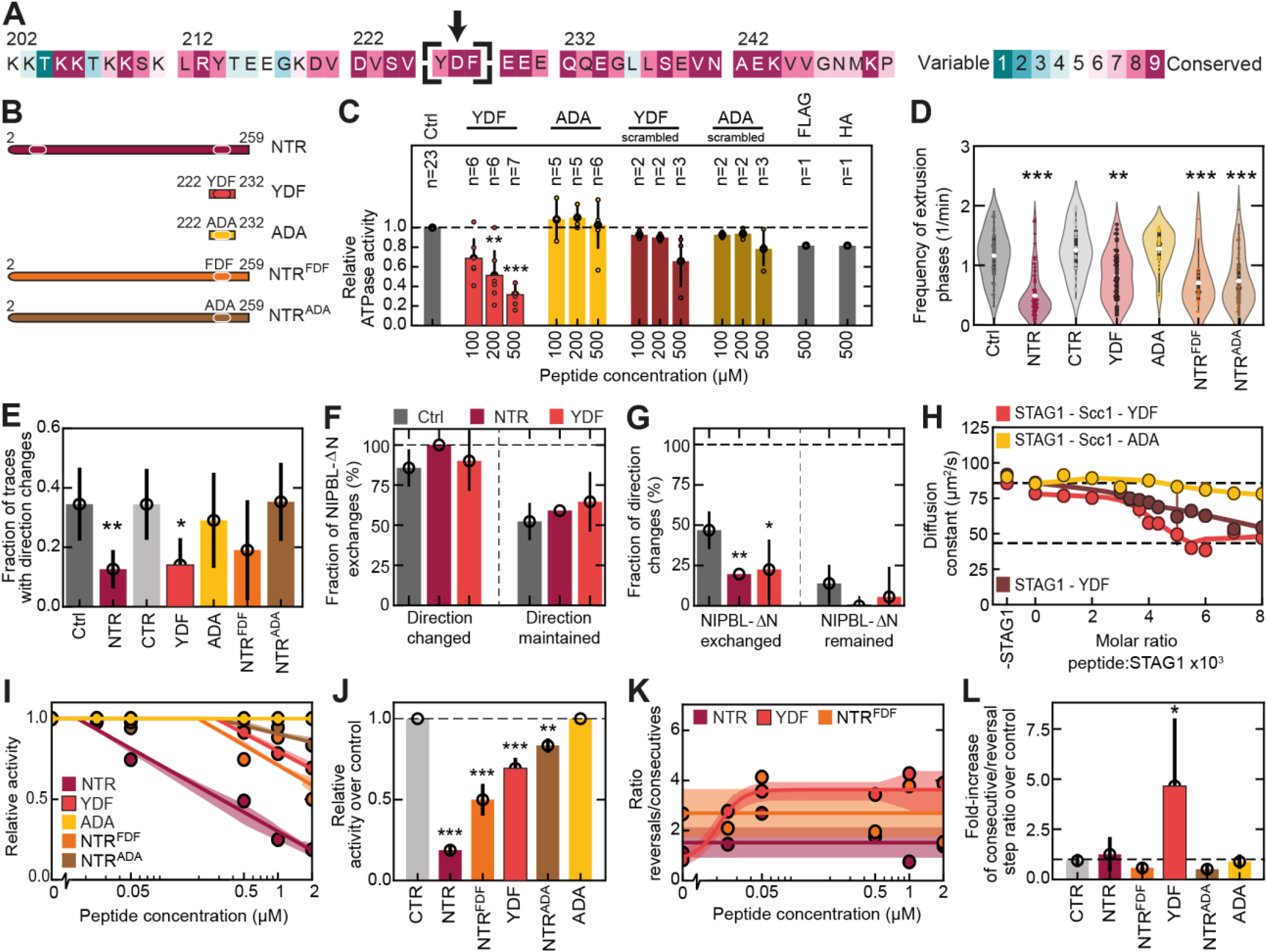
CTCF’s YDF motif impairs cohesin’s LE activity and renders LE unidirectional. (**A**) Conservation annotation of the human CTCF sequence surrounding the conserved YDF motif. (**B**) Illustration of CTCF fragments used. (**C**) Normalised ATPase activity of cohesin for various concentration of CTCF peptide. Scrambled peptides contain the same amino acids as the non-scrambled version but in random order. Statistical significance was assessed by an ANOVA with Holm-Sidak’s multiple comparison test (**: p < 0.01; ***: p < 0.001). (**D**) Violin plot of the frequency with which extrusion phases occur in the presence of 2 μM of the indicated CTCF fragment as in Fig. 1D. N = 45, 91, 40, 19, 49, 52, 25 DNA molecules from left to right. (**E**) Fraction of traces that show direction changes in the absence (control) or the presence of 2 μM of the indicated CTCF fragment as in Fig. 1E. N = 58, 103, 57, 21, 51, 57, 32 DNA molecules from left to right. (**F**) Fraction of cohesin molecules for which NIPBL-ΔN exchanged in between direction changes (left; N = 35, 23, 27 from left to right) and the fraction of cohesin molecules on which NIPBL-ΔN exchanged without direction change between successive extrusion phases (right; N = 65, 80, 64). Error bars denote the binomial 95% confidence interval. Statistical significance was assessed using a Chi-square test. (**G**) Fraction of direction changes between successive extrusion phases during which NIPBL-ΔN exchanged (N = 64, 65, 59) and the fraction of direction changes between successive extrusion phases during which NIPBL-ΔN remained (N = 36, 38, 32). Statistical significance was assessed using a Chi-square test (*: p < 0.05; **: p < 0.01). (**H**) Diffusion constant of DNA-STAG1^+/-Scc1^- YDF and DNA-STAG1^Scc1^-ADA complexes versus the molar ratio of peptide to STAG1 derived from a monoexponential fit to FCS curves. Upper and lower dashed lines denote the theoretical diffusion constant of freely diffusing DNA and the DNA-STAG1^Scc1^-YDF (or -ADA) complex, respectively. Experiments were conducted with two biological replicates and 4–9 technical replicates. Data points and error bars represent the mean ± SD of these replicates, while solid lines display a LOWESS-smoothed trend of the means. (**I**) The relative stepping activity of cohesin, compared to the control ([peptide] = 0 μM), versus peptide concentration as in Fig. 1I. The data corresponds to the inverse of the red bar height of panels Fig. 1H, Fig. S3F-I. Fits serve as a guide for the eye (Methods). Lines denote fits and the shaded error bars denote the interpolated SD of the individual data points. (**J**) The relative stepping activity, compared to the control, for 2 μM of the indicated peptides. Error bars denote the standard deviation from repeated experiments. Statistical significance was assessed by a one-sided Welch’s t-test (**: p < 0.01; ***: p < 0.001). (**K**) The ratio of reversal and consecutive steps with increasing concentrations of the indicated peptides from data in Fig. 1H, Fig. S3F-I. See also Fig. S3J. (**L**) The ratio of reversal and consecutive steps for 2 μM of the indicated peptides as in (J).

Unexpectedly, upon analysing LE direction changes, we found that the YDF motif alone caused cohesin to extrude unidirectionally, mimicking the effect of the full NTR (Fig. 2E). While the Y226F mutation produced an intermediate (not statistically significant) effect, the Y226A/F228A mutation and the ADA peptide completely abolished the unidirectional bias. These results indicate that the YDF motif alone drives cohesin’s unidirectional extrusion. Direction changes in cohesin-mediated LE has been demonstrated to require NIPBL exchange (*15*). To test whether the YDF motif affects cohesin’s directionality by preventing NIPBL exchange, we conducted single-molecule fluorescence experiments with an equimolar mix of NIPBL-ΔN molecules that were fluorescently labeled in two colors (*15*) (Methods). The presence of the NTR or YDF peptides was found to not alter the residence time of NIPBL-ΔN on cohesin (Fig. S3C), ruling out a slower NIPBL-ΔN exchange as the cause of unidirectional extrusion. We then correlated NIPBL presence, absence, and exchange with extrusion direction switches. Both in the absence and presence of the NTR or the YDF motif, almost all direction changes required an NIPBL-ΔN exchange (Fig. 2F). Similarly, NIPBL exchanged in ∼50% of cases when the extrusion direction was maintained between successive extrusion phases (Fig. 2F). In control experiments, roughly 50% of the exchanges of NIPBL-ΔN coincided with a direction switch (Fig. 2G). However, in the presence of the NTR or the YDF motif, we observed that 2.5-fold fewer NIPBL-ΔN exchange events led to direction changes (Fig. 2G), suggesting that the NTR and YDF motif prevent direction changes through a mechanism unrelated to NIPBL exchange dynamics.

Directionality switching in SMC-mediated loop extrusion is thought to involve DNA strand exchange between the extruding and anchoring sides of the SMC complex (*6*, *15*, *56*). Data that showed that yeast condensin, a strictly unidirectional extruder, extrudes bidirectionally upon deletion of its anchoring Ycg1 subunit (*56*) led us to hypothesize that the YDF motif might inhibit DNA strand exchange by enhancing the DNA-binding affinity of STAG1. To test this, we conducted Fluorescence Correlation Spectroscopy (FCS) experiments that measure the diffusion constant of fluorescently labeled DNA in the presence of purified STAG1, Scc1^251-420^, and YDF peptide (Methods). Increasing concentrations of the YDF peptide caused the DNA’s diffusion constant to shift from that of freely diffusing molecules to STAG1-bound complexes, indicating an increased DNA affinity (Fig. 2H). This effect was partially dependent on Scc1 and largely absent with the ADA peptide (Fig. 2H).

MT experiments with increasing YDF peptide concentrations showed a ∼30% decrease in cohesin’s stepping activity at 2 μM (Fig. 2I,J; Fig. S3D-F). While consecutive steps decreased (Fig. S3F,K), the fraction of reversals remained constant up to 1 µM YDF peptide (Fig. S3F,L), resulting in an increased ratio of reversals to consecutive steps (Fig. 2K,L). These effects were absent with the ADA peptide (Fig. S2I,J,L; Fig. S3G,J-L). YDF-mutated NTR variants (NTR^FDF^ and NTR^ADA^) also reduced cohesin activity like the wild-type NTR but had progressively weaker effects, which were detected only at higher concentrations (Fig. 2I,J; Fig. S3H,I). Similar to the wild-type NTR (Fig. S1M), NTR^FDF^ and NTR^ADA^ did not affect the fraction of reversals and consecutive steps (Fig. 2K,L; Fig. S3J-L). The reduced effect of these mutations on cohesin-mediated LE is likely due to lower binding affinity of the YDF mutations compared to the wild-type YDF motif to the CES (*33*), as observed previously for similar motifs in WAPL (*35*). Cohesin-mediated LE is essential for the formation of TADs as observed in Hi-C matrices (*12*, *14*, *16*–*20*). A prevailing hypothesis suggests that cohesin stalls at a CTCF site on the encountering side and extrudes toward the opposite side until blocked by a second CTCF. However, single-molecule experiments revealed that cohesin rarely reverses direction after encountering CTCF (*26*) and fully looped TADs are rarely observed *in vivo* (*57*, *58*). Our findings indicate that the CTCF NTR inhibits cohesin, and the YDF motif prevents cohesin from switching directions and extruding away from the encountered CTCF. To explore whether cohesin stalling, as opposed to direction reversal, can account for the patterns seen in Hi-C, we performed 3D polymer simulations (Methods). These simulations indicate that experimentally observed Hi-C maps are well explained by stalled cohesin which does not extrude away from encountered CTCF sites (Supplementary Text; Fig. S4).

In summary, the YDF motif biases cohesin towards unidirectional extrusion by strengthening the DNA affinity of STAG1 to DNA (Fig. 2H), likely preventing DNA strand exchanges. The wild-type NTR reduces cohesin’s activity most significantly, followed by NTR^FDF^ and the YDF motif alone. The effect of the YDF peptide is dependent on the YDF motif, since mutation to ADA alleviates this inhibition (Fig. S2I,J). While the YDF motif alone increases the fraction of reversals relative to consecutive steps, the full NTR reduces both (Fig. 2K,L; Fig. S3K,L). We conclude that CTCF’s NTR regulates cohesin activity through mechanisms that extend beyond the YDF motif alone.

## The KTYQR motif in the CTCF NTR inhibits cohesin-mediated loop extrusion

Nora *et al.* (*41*) identified CTCF regions that impact TAD insulation, including the N-terminal region NTR^1-89^, an RNA-binding zinc finger domain (NTR^264-288^), and YDF motif. We evaluated NTR^1-89^, which contains a conserved KTYQR motif at residues 23-27 (Fig. 3A; Fig. S2B) that is known to bind PDS5A (*41*). We tested the effect of a fragment containing the KTYQR motif (residues 16-31; ‘KTYQR peptide’; Fig. 3B; Table S1) on cohesin’s LE dynamics, and found that it strongly affected it: The KTYQR peptide reduced cohesin’s ATPase activity by ∼60%, similarly to the YDF peptide (Fig. 3C), and it halved the frequency of active LE phases (Fig. 3D), comparable to the NTR^FDF^ and NTR^ADA^ mutants, although it did not affect the LE rate (Fig. S5B) or directionality (Fig. 3E).

**Figure 3.**
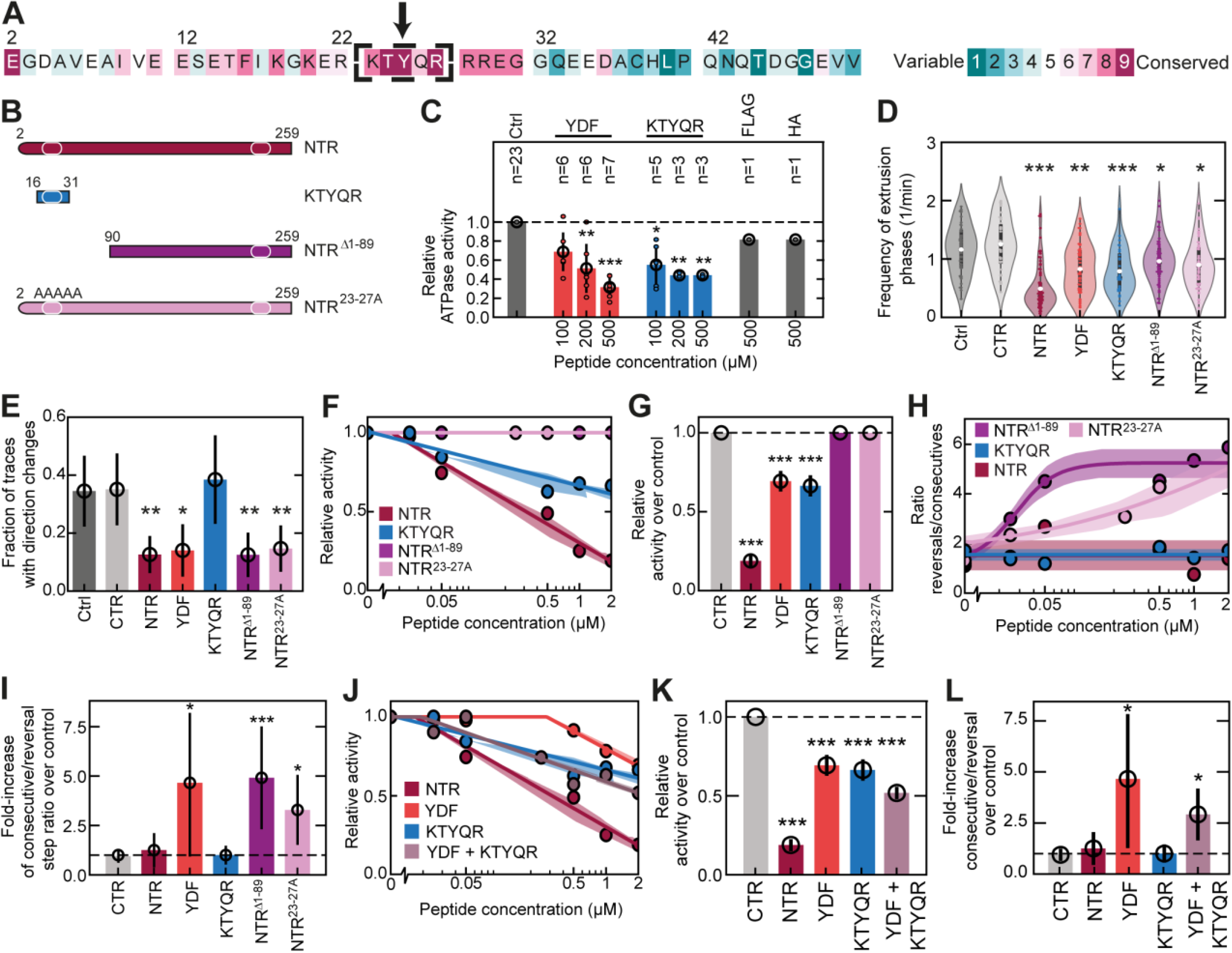
CTCF’s KTYQR and YDF motifs inhibits LE by cohesin. (**A**) Conservation annotation of the human CTCF sequence surrounding the KTYQR motif. (**B**) Illustration of the short KTYQR peptide as well as CTCF NTR and the corresponding truncation and mutation. (**C**) Normalised ATPase activity of cohesin in presence of the indicated concentration of CTCF peptide. (**D**) Violin plot of the frequency with which extrusion phases occur in the presence of 2 μM of various CTCF fragment as in Fig. 1D and Fig. 2D. N = 45, 40, 91, 19, 29, 62, 58 DNA molecules from left to right. (**E**) Fraction of traces that show direction changes in the absence (control) or the presence of 2 μM of the indicated CTCF fragment. N = 58, 57, 103, 21, 39, 72, 75 DNA molecules from left to right. (**F**) Relative stepping activity of cohesin, compared to the control ([peptide] = 0 μM), for increasing concentrations of the indicated peptides as in Fig. 1I and Fig. 2I. Data corresponds to the red bar height of panels Fig. 1H, Fig. S5E-G. Fits serve as a guide for the eye (Methods). Lines denote the fit and the shaded error bars denote the interpolated SD of individual data points. (**G**) Relative stepping activity for various peptides at 2 μM, compared to control. Error bars denote the standard deviation from repeated experiments. Statistical significance was assessed by a one-sided Welch’s t-test (**: p < 0.01; ***: p < 0.001). (**H**) The ratio of reversal and consecutive steps with increasing concentrations of the indicated peptides from data in Fig. 1H, Fig. S5E-G. (**I**) Ratio of reversal and consecutive steps for 2 μM of the indicated peptides as in (G). (**J**) Relative stepping activity of cohesin, compared to the control ([peptide] = 0 μM), versus concentration of the indicated peptides as in Fig. 1I, Fig. 2I, and Fig. 3F. The data corresponds to the inverse of the red bar height of panels Fig. 1H, Fig. S3F, Fig. S5E, and Fig. S5J. (**K**) Ratio of reversal and consecutive steps with increasing concentrations of the indicated peptides from data as in (J). (**L**) The ratio of reversal and consecutive steps for 2 μM of the indicated peptides for the same data as in (J).

To probe the effects of the NTR without the KTYQR motif on cohesin, we expressed and purified an N-terminal truncation of the NTR (NTR^Δ1-89^) as well as an alanine-substituted mutant (NTR^23-27A^; Fig. 3B, Fig. S5A; Table S1). Both variants caused only a modest reduction in LE frequency (20% and 25% lower median values, respectively; Fig. 3D), suggesting that the KTYQR motif is the major determinant. While they did not affect the LE rate (Fig. S5B), both variants converted cohesin into a unidirectional extruder (Fig. 3E), consistent with the major role of the YDF motif in setting the directionality.

In MT experiments, the KTYQR peptide significantly reduced cohesin’s stepping activity (Fig. 3F,G; Fig. S3C-E), achieving effects similar to 2 μM YDF peptide but the inhibiting effect appears at lower concentrations (Fig. 2I; Fig. 3F,G). By contrast, the N-terminal truncation NTR^Δ1-89^ and the alanine-substituted mutant NTR^23-27A^ showed no noticeable reduction of LE activity (Fig. 3G). Unlike YDF, the KTYQR peptide reduced both consecutive and reversal steps equally (Fig. 3H,I; Fig. S5H,I). NTR variants lacking KTYQR (NTR^Δ1-89^ and NTR^23-27A^) did not affect the overall stepping activity (Fig. 3F,G; Fig. S5F,G), but increased the occurrence of reversals while reducing consecutive steps (Fig. S5H,I), raising the ratio of reversal to consecutive steps (Fig. 3H,I). These findings suggest that the absence of the KTYQR motif amplifies the YDF motif’s effect in the NTR, favoring reversal steps at the expense of productive extrusion.

We thus conclude that the YDF and KTYQR motifs of the NTR of CTCF distinctly affect cohesin-mediated LE action. Both motifs suppress cohesin’s overall activity (Fig. 3G), but they exhibit different effects on LE dynamics. The YDF motif preferentially inhibits consecutive LE steps at lower concentrations, while KTYQR reduces both consecutive and reversal steps without altering the extrusion directionality. Together, the YDF and KTYQR motifs synergistically reduce cohesin’s activity.

Finally, we examined whether the combined effect of the YDF and KTYQR peptide fragments could replicate the effect of the full NTR on cohesin-mediated LE. In MT experiments, adding both peptides reduced cohesin’s stepping activity significantly more than either peptide alone, but still less than their additive individual effects or the full NTR (Fig. 3J,K). Notably, the combination of YDF and KTYQR peptides impaired cohesin’s ability to take consecutive LE steps to a degree similar to the full NTR and surpassing the effect the YDF peptide has alone (Fig. S5L). However, their effect on reversal steps was intermediate between that of the individual peptides and the full NTR (Fig. S5M), reducing the effect on the ratio of consecutive to reversal steps (Fig. SK; Fig 3L). These results suggest that the full NTR exerts additional suppression on reversal steps, whereas YDF and KTYQR motifs together can account for the NTR’s inhibition of consecutive LE steps and thus suffice to account for CTCF’s ability to stall cohesin.

## CTCF and cohesin are predicted to interact through KTYQR binding to STAG1 and YDF binding to STAG1 and hinge

The YDF motif binds to cohesin at the CES, a surface formed by STAG1/2 and Scc1(*33*–*39*). To explore if and how other CTCF NTR regions, such as the KTYQR peptide, bind to and affect cohesin, we used AlphaFold-Multimer (*59*, *60*) to predict putative binding sites across cohesin subunits and CTCF NTR fragments. Cohesin was segmented into subunits (NIPBL-Mau2, STAG1/2, ATPase heads, and hinge regions with Scc1 segments; Methods; Table S2; Table S3) and CTCF was divided into three overlapping fragments (#1: 1-100, including KTYQR; #2: 90-190; #3 180-264, including YDF; Table S2).

AlphaFold predicted interactions between STAG1/2 and CTCF fragments #1 and #3 (Fig. 4A; Fig. S6A-C) and between the Smc hinge and fragment #3 (Fig. S6D,E). Consistent with previous experimental finding, interactions between STAG1/2 and fragment #3 were mediated by the CES and the YDF motif (*33*). A new finding, however, is that in most predictions (16/25 or 25/25 for STAG1 and STAG2, respectively), the KTYQR motif was found to bind to a conserved surface, N-terminally of the CES (Fig. 4A and Fig. S6A,B). As AlphaFold does not quantitate binding affinities, the KTYQR motif may bind transiently or with low affinity, making it challenging to detect in bulk (*33*) or with *in vivo* binding assays (*41*). To experimentally test the AlphaFold prediction, we performed FCS using a fluorescently labeled KTYQR peptide and purified STAG1 (Methods). Increasing the molar ratio of STAG1 over the KTYQR peptide showed an increasing fraction of KTYQR peptide with a diffusion constant corresponding to a STAG1^KTYQR^ complex (Methods; Fig. 4B), indicating that KTYQR indeed binds to STAG1.

**Figure 4.**
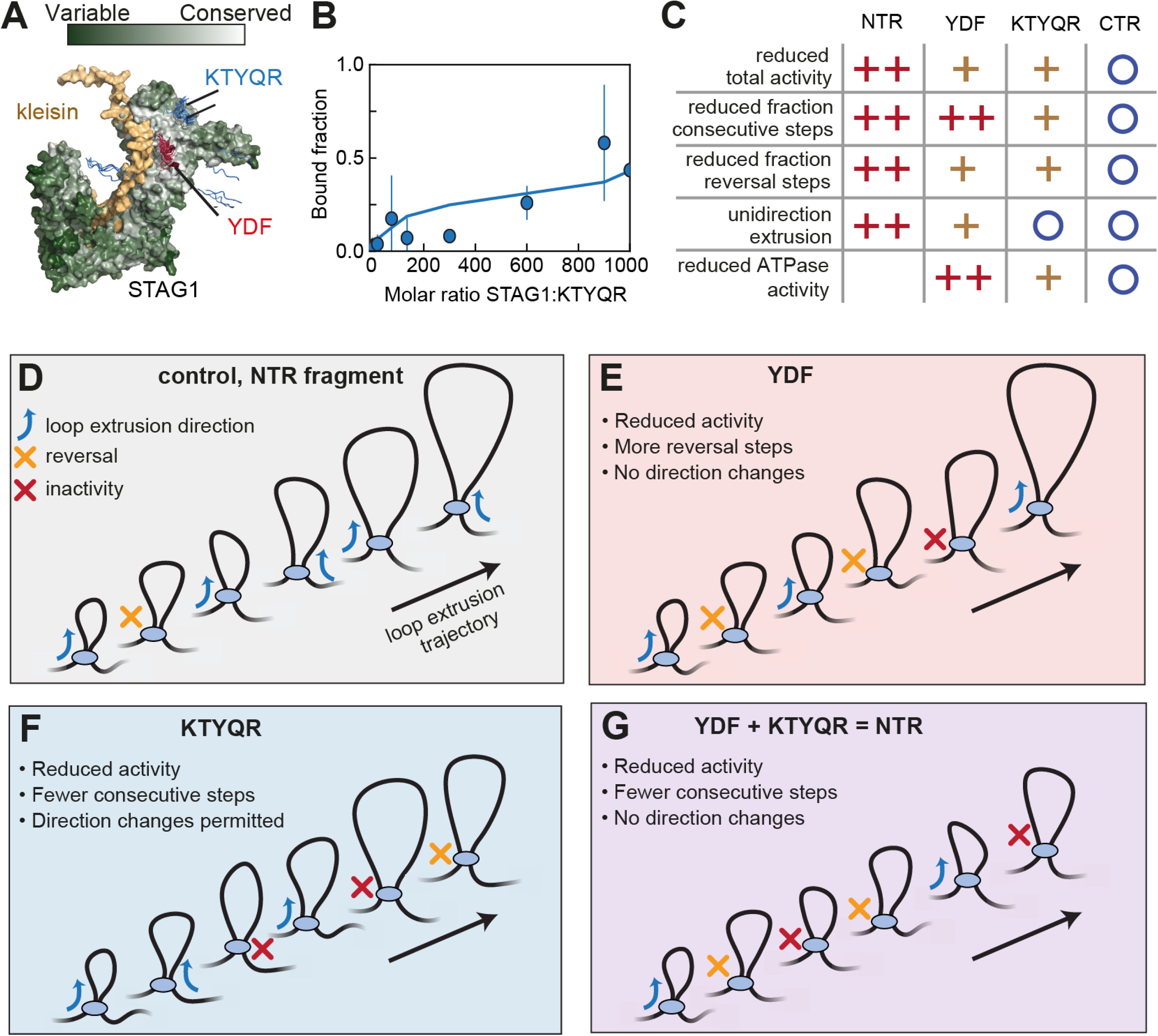
CTCF’s YDF and KTYQR account for most of CTCF’s effect on cohesin. (**A**) AlphaFold prediction of the STAG1^Scc1^ complex and 25 predictions of the placement of the YDF and KTYYQR peptides. STAG1 was colored according to the conservation score (Methods). (**B**) Fraction of fluorescently labeled KTYQR peptide bound to STAG1 versus the molar ratio of STAG1 to KTYQR peptide (Methods). Experiments were conducted with two biological replicates and 4–9 technical replicates. Data points and error bars represent the mean ± SD of these replicates, while solid lines display a Savitzky-Golay-smoothed (window of 7 points, order 1) trend of the means. (**C**) Overview of the effect of the NTR, YDF motif, KTYQR motif, and CTR on various characteristics of LE by cohesin (O: no effect; +: intermediate effect; ++: strong effect). (**D-G**) Illustrations of the effect of the YDF, KTYQR, and their combination on DNA loop extrusion by cohesin.

AlphaFold also predicted an interaction between the YDF motif and the Smc hinge in 18/25 predictions (Fig. S6D,E). This interaction might regulate LE by cohesin and hinge opening to modulate sister chromatid cohesion (*61*–*64*). Supporting this, Nagasaka *et al*. (*65*) identified a separation-of-function mutant (SMC1^K540D/K637D/K648D^) that retained LE but is defective in cohesion, similar to other hinge mutants previously identified in yeast (*64*). Curiously, this mutant was also reduced in forming CTCF-anchored loops but not TADs *in vivo* (*65*). It is tempting to speculate that an interaction between the cohesin hinge and YDF inhibits LE and thus contributes to the formation of CTCF-anchored loops. This might also modulate cohesin’s tendency to undergo unsuccessful LE step cycles (reversals) due to changes in the DNA binding affinity of the Smc hinge, which is involved in LE (*66*).

Together, the results suggest that the KTYQR motif inhibits cohesin by binding to STAG1/2, while the YDF motif may regulate LE through interactions with the CES and the hinge. Possibly, the YDF-hinge interaction may also regulate cohesion.

## Concluding remarks

Our study provides mechanistic insights into how CTCF inhibits cohesin-mediated LE (*12*, *14*, *16*–*18*, *20*, *26*, *27*, *67*). Our data clearly show that the NTR rather than the CTR of CTCF is responsible for the interaction with cohesin. Nevertheless, prior *in vitro* studies observed that cohesin was also able to halt LE when approached from the C-terminal side of CTCF (*26*). Our findings suggest that in such cases, cohesin may interact with CTCF’s N-terminus. Supporting this, the residence times of cohesin and CTCF are similar between N- and C-terminal encounters (*26*).

While previous studies established CTCF’s crucial role in stalling cohesin, implying YDF-CES binding of STAG1/2 *in vivo* (33), the molecular details remained unclear. To reveal the underlying mechanism, we studied the effects of the CTCF NTR on cohesin-mediated LE *in vitro* and revealed the individual contributions of the YDF and KTYQR motifs therein (Fig. 4C). The YDF motif reduces cohesin’s activity by decreasing ATPase activity, reducing consecutive loop-extrusion steps, and driving the extrusion unidirectional by increasing STAG1’s DNA-binding affinity, which presumably prevents DNA strand exchange (Fig. 2; Fig. 4D,F) (*15*, *56*, *68*). While the KTYQR motif also reduces ATPase activity, it halts the overall LE stepping activity, in contrast to YDF (Fig. 3F,G; Fig. 4E). Combined, these two motifs fully suppress cohesin’s ability to take consecutive LE steps (Fig. 3J-L; Fig. 4G).

Our study reveals that CTCF’s N-terminal region modulates cohesin’s LE activity and directionality through its YDF and KTYQR motifs via distinct yet synergistic mechanisms. Future studies may aim to capture the temporal sequence of conformational changes in cohesin leading to LE to further elucidate the particular binding states within the LE cycle. Such studies could also address in more detail how modulating STAG1’s DNA binding affinity leads to a change in both the cohesin’s ATPase activity and the LE initiation efficiency.

## Acknowledgments

We thank Brian Analikwu, Allard Katan, and Miloš Tišma for discussions; Mathias Madalinski for purifying Atto550- and JF646-HaloTag Ligands as well as for synthesising fluorescently labeled KTYQR peptide; Melanie Panarotto for purifying STAG1; Marcel van den Broek for help with setting up AlphaFold2. The authors acknowledge the use of computational resources of the DelftBlue supercomputer, provided by Delft High Performance Computing Centre (https://www.tudelft.nl/dhpc). J.-M.P. is also an adjunct professor at the Medical University of Vienna.

## Funding

ERC Advanced Grant 883684 (DNA looping) (CD)

BaSyC program (CD)

EVOLF program (CD) Boehringer Ingelheim (JMP)

Austrian Research Promotion Agency Headquarter grant FFG-FO999902549 (JMP)

European Research Council under the European Union’s Horizon 2020 research and innovation program GA No 101020558 (JMP)

Vienna Science and Technology Fund grant LS19-029 (JMP)

## Author contributions

Conceptualisation: RB, RJ, JvdT, CD

Data curation: RB, RJ, LM, GL

Formal analysis: RB, RJ, LM, GL

Methodology: RB, RJ, LM, JvdT, GL, ID, JMP, CD

Investigation: RB, RJ, LM, GL

Visualisation: RB, RJ, LM

Funding acquisition: JMP, CD

Project administration: RB, RJ, JMP, CD

Resources: RB, RJ, LM, GL, EvdS, AvdB, ID

Software: RB, RJ, LM

Supervision: JMP, CD

Validation: RB, RJ, LM, GL

Writing – original draft: RB, RJ, LM

Writing – review & editing: RB, RJ, LM, JvdT, GL, EvdS, AvsB, IF, JMP, CD

## Data and materials availability

Requests for materials, data, and software generated in this study should be directed to the lead contact.

## Supplementary Materials

Materials and Methods

Supplementary Text

Figs. S1 to S6

Tables S1 to S3

References (*69*–88)

---

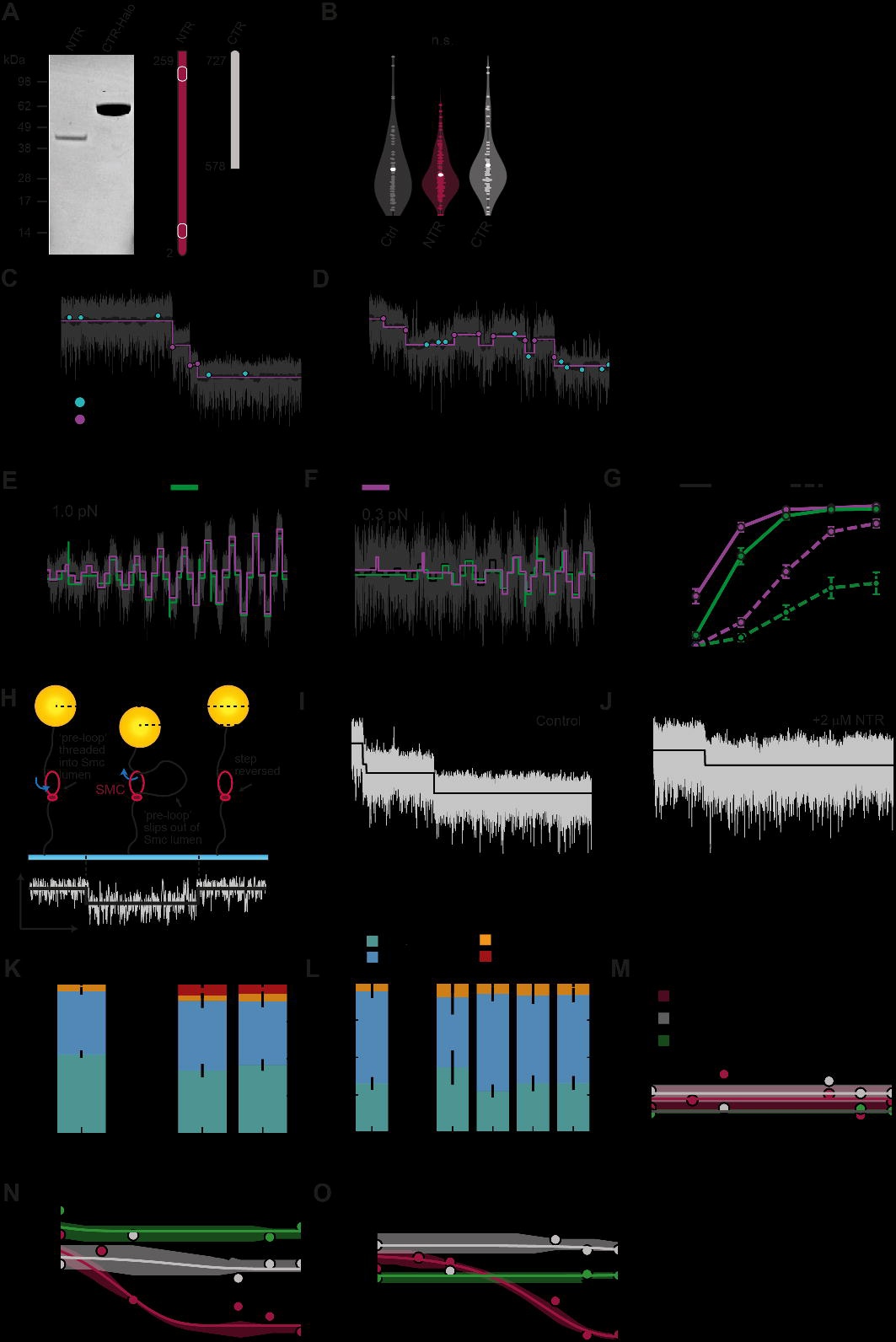

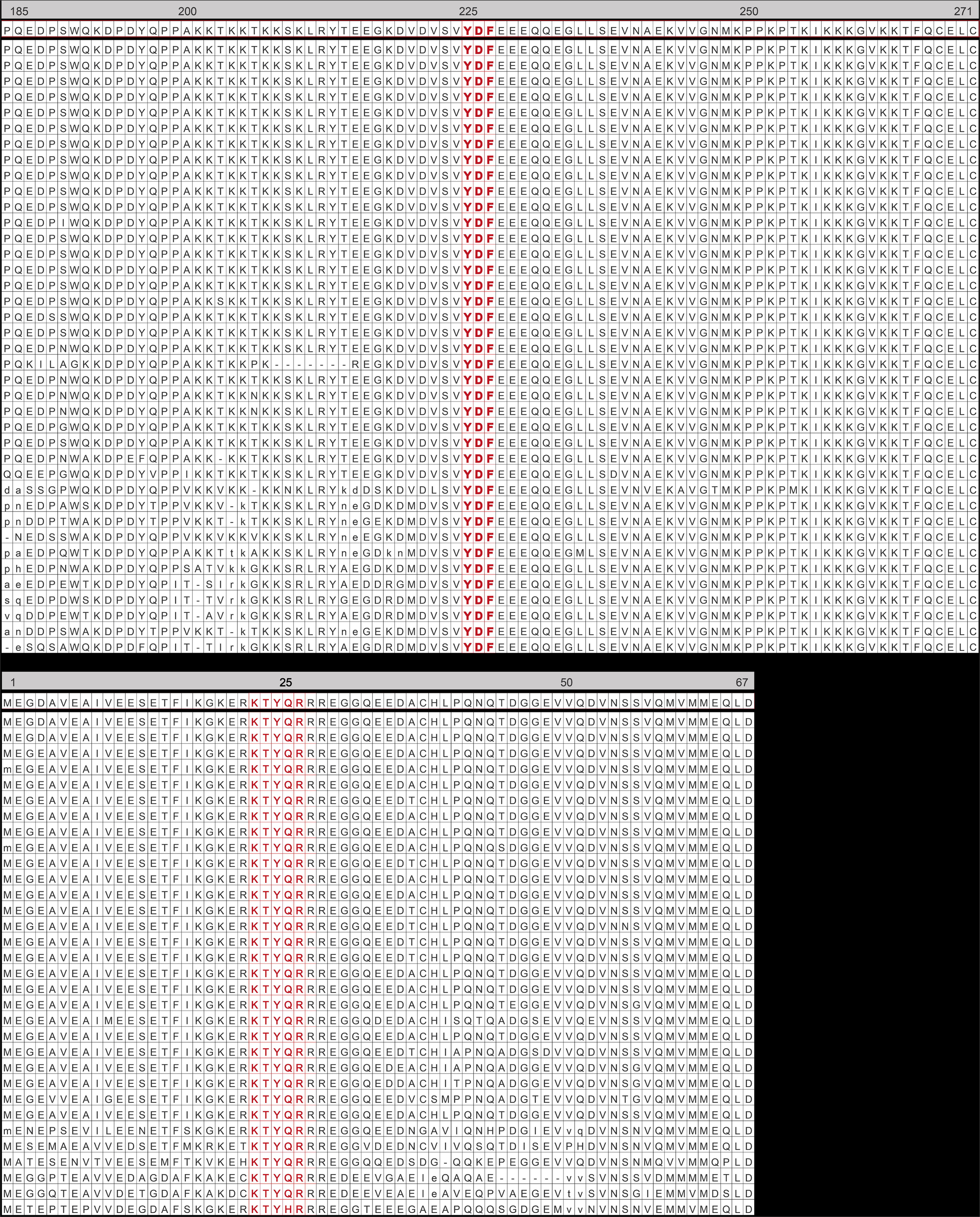

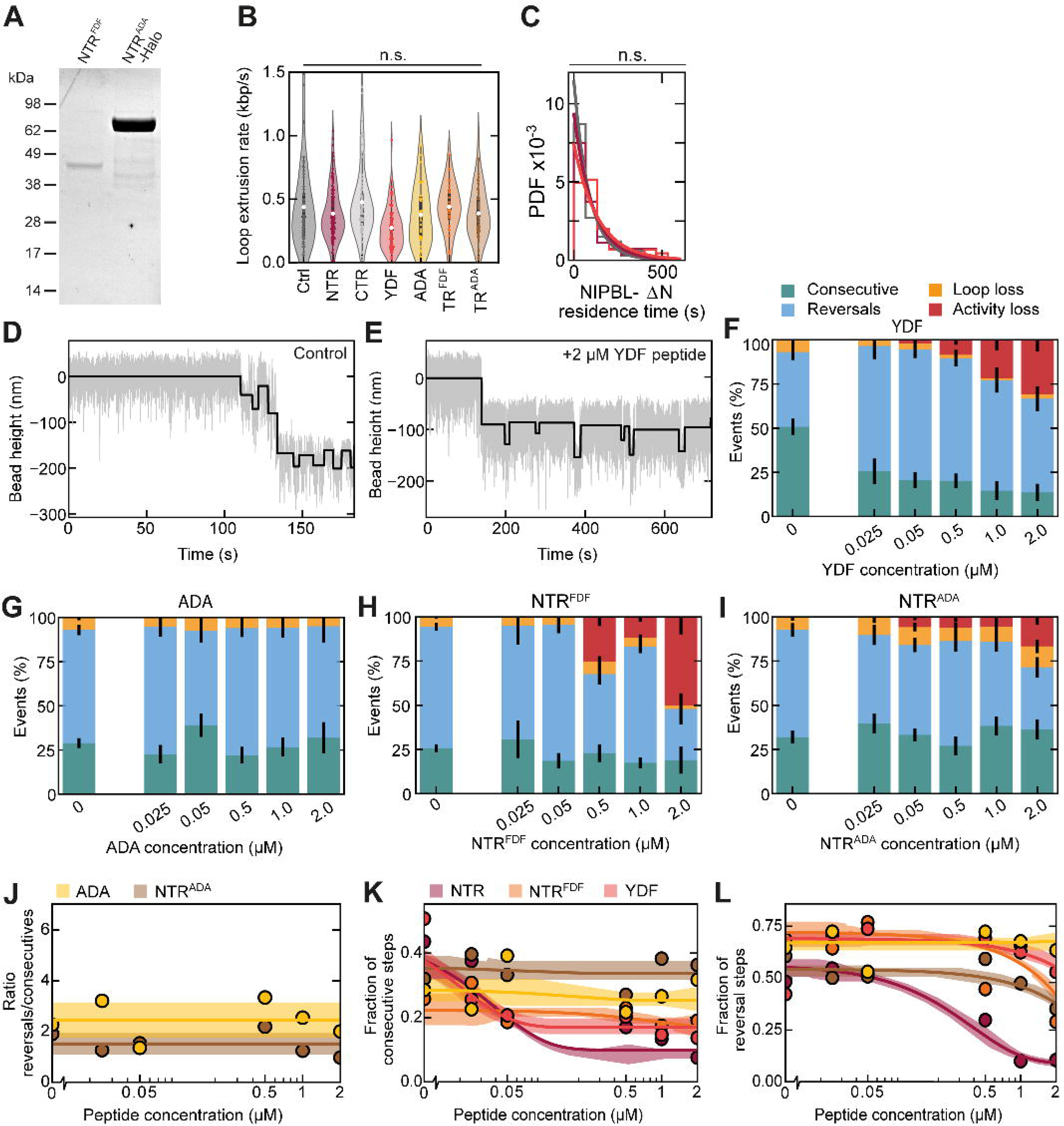

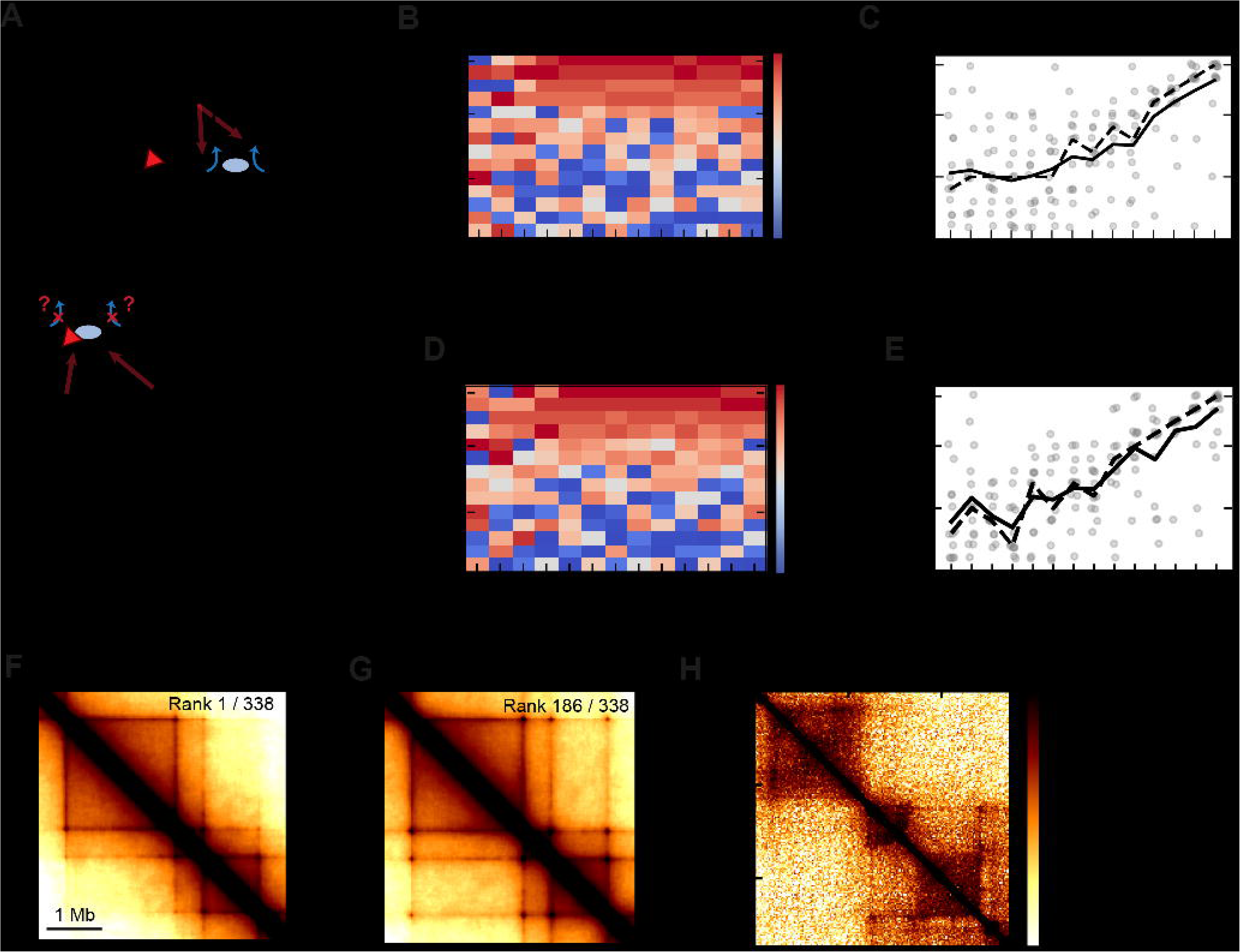

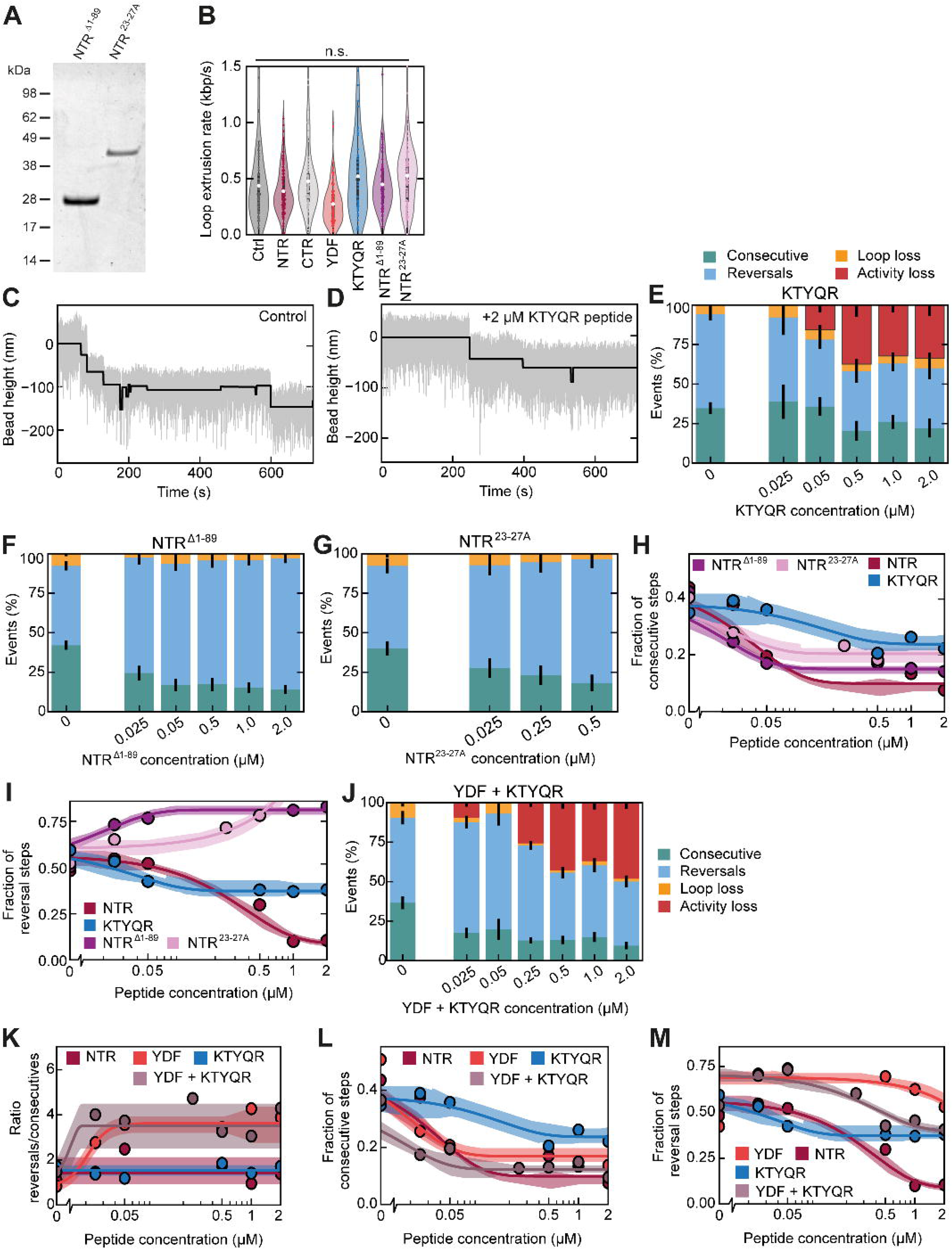

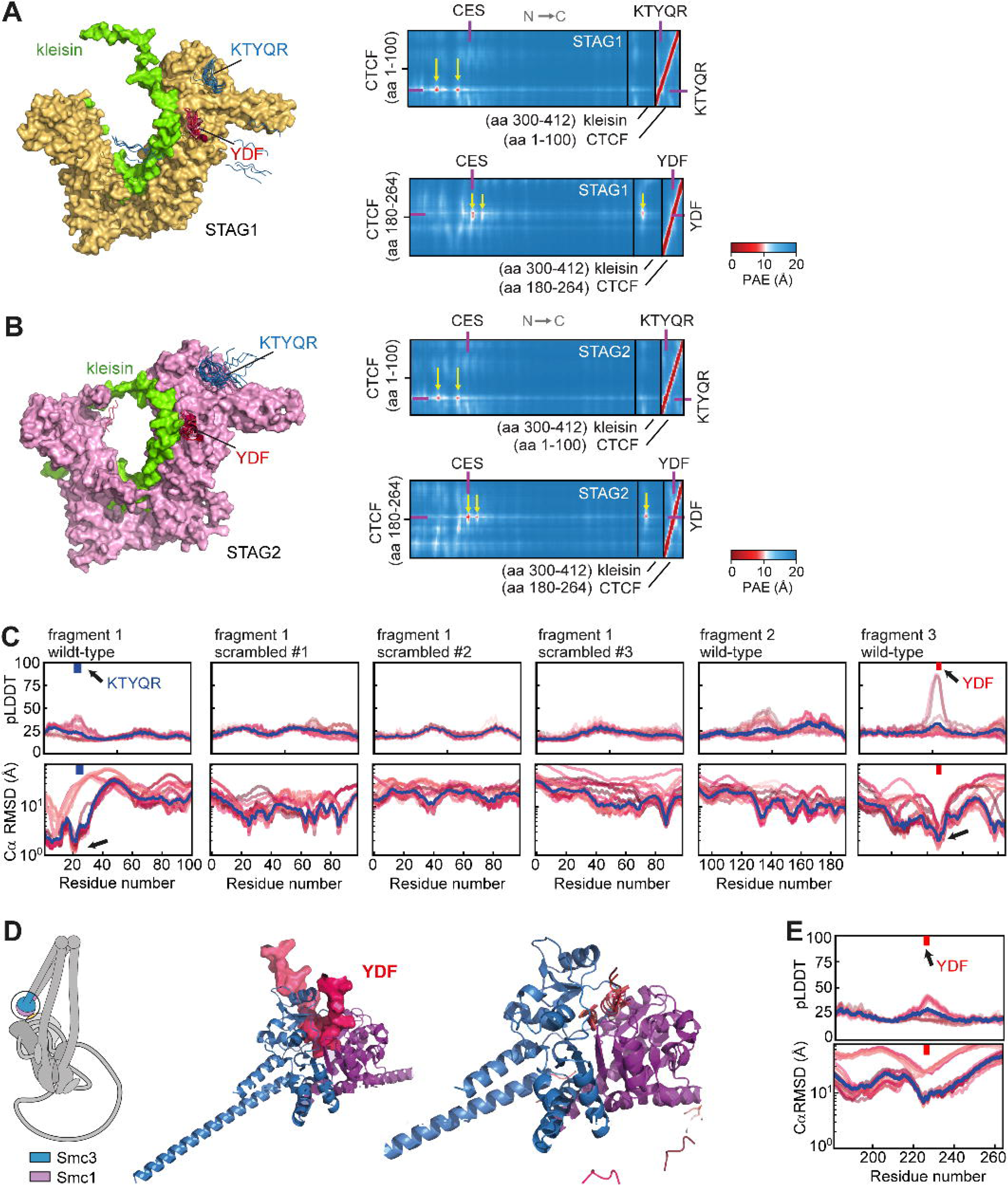

