## Supplementary Information for "Two CTCF motifs impede cohesin-mediated DNA loop extrusion"

### Materials and Methods

Human cohesin, NIPBL-Mau2, STAG1, and NIPBL-ΔN expression and purification

Wild-type human cohesin, NIPBL-Mau2, and STAG1 were expressed in and purified from *Sf9* insect cells as described previously (*3*). NIPBL-ΔN was expressed, purified, split in half, and fluorescently labeled with Atto550 and JF646, respectively, as described previously (*15*). As before (*15*), protein and fluorophore concentrations were measured at 280 nm for NIPBL-ΔN and at 550/646 nm for A550/JF646, yielding a fraction of 86 ± 4 % labeled NIPBL-ΔN-A550 (N = 7) and 85 ± 5 % labeled NIPBL-ΔN-JF646 (N = 5) molecules. FCS was performed on samples of 10 nM NIPBL-ΔN-A550/JF646 in a buffer containing 40 mM Tris-HCl pH 7.5, 50 mM NaCl, 2.5 mM MgCl2, 1 mM DTT at room temperature. The resulting autocorrelation functions were well described by a fit with a single component, indicating that no free fluorophores were present.

Yeast condensin expression and purification

Wild-type yeast condensin was expressed and purified from *Saccharomyces cerevisiae* as described previously (*4*).

Expression and purification of CTCF fragments

Various fragments of human CTCF (Uniprot #P49711, see Table S1) were cloned into a pMAL-C5X derived plasmid, resulting in MBP fusions with a flexible linker and a 3C protease site at the fusion point (MBP-NSSSNNNNNNNNNNLGIEGRISHMSMGGRDIVDGSEF-LEVLFQ ↓GP-CTCF). Plasmids were transformed into *E. coli* ER2566 cells (New England Biolabs, fhuA2 lacZ::T7 gene1 [lon] ompT gal sulA11 R(mcr73::miniTn10--TetS)2 [dcm] R(zgb-210::Tn10--TetS) endA1 Δ(mcrCmrr)114::IS10) containing a Rosetta®-2 plasmid (Novagen), and selected on LB media containing 100 µg/ml ampicillin and 34 µg/ml chloramphenicol. For expression, overnight pre-cultures were diluted 1:100 into 1 liter prewarmed LB with fresh antibiotics, cultures were grown in baffled flasks shaking at 200 rpm at 37°C until the OD600 reached ~0.6, and expression was induced by the addition of IPTG to a final concentration of 1 mM. After 3 hours of induction, the cells were harvested by centrifugation (8 min, 4500 rpm, JLA8.1000 rotor), washed in 50 ml PBS, and resuspended in buffer A (50 mM Tris-HCl pH 7.5, 300 mM NaCl). Cells were lysed using a French Press (Constant Systems) at 20 kpsi, 4°C and unbroken cells, debris and aggregates were pelleted in a Ti45 rotor (30 min, 40.000 rpm, 4°C). The lysate was applied to 2 ml prewashed amylose resin (NEB #E8021) and incubated for one hour while rotating at 4°C. Subsequently, the resin was washed with 50 ml of buffer A and finally, CTCF fragments were eluted with 15 ml of buffer A supplemented with 1 mM β-mercaptoethanol and homemade 3C protease. CTCF fragments were concentrated using a Vivaspin centrifugal concentrator (10 kDa cut-off) and further purified by size exclusion chromatography (SEC) on a Superdex 200 Increase 10/300 column pre-equilibrated with buffer B (50 mM Tris-HCl pH 7.5, 150 mM NaCl, 5% (w/v) glycerol, 0.05 mM TCEP).

To purify the CTCF CTR (10xHis-CTCF(578-727)-TEV-Halo) and NTR$^{ADA}$ (10xHis-CTCF(2-259)-TEV-Halo), baculovirus-infected insect *Sf9* cell pellets from cultures supplemented with 0.1 mM $ZnCl_2$ were lysed by Dounce homogenisation and resuspended in CTCF buffer (35 mM Hepes pH 7.5, 350 mM NaCl, 5% glycerol) supplemented with 0.1 mM $ZnCl_2$, 0.05% Tween-20, 5 mM imidazole, 1 mM PMSF, EDTA-free cOmplete tablet (1 per 50 ml) (Roche, 11873580001), 3 mM betamercaptoethanol, 10 µg/ml aprotinin, 2 mM benzamidine and benzonase. The lysate was cleared by centrifugation at 18,000 g for 45 min at 4°C. The soluble fraction was incubated with NiNTA agarose (Qiagen, 30230) for 90 min at 4°C and washed first with CTCF buffer supplemented with 0.1 mM $ZnCl_2$, 0.01 % Tween-20, 35 mM imidazole, 1 mM PMSF and 2 mM benzamidine and then with CTCF buffer supplemented with 35 mM imidazole only. Protein was eluted with CTCF buffer supplemented with 300 mM imidazole. The eluate was concentrated using a Vivaspin 30 kDa MWCO concentrator (Sartorius) and buffer was exchanged using a Zeba 7 kDa MWCO desalting column equilibrated in CTCF buffer. Protein was then flash-frozen and stored at -80°C.

In addition to these longer protein fragments, smaller fragments were synthesised in-house or ordered from Genscript (Table S1). Peptides were synthesised on a Liberty Blue peptide synthesiser (CEM) using standard Fmoc chemistry. For each amino acid cycle, a 4 min coupling with DIC/Oxyma was performed. Peptides were purified on a Phenomenex Luna C18(2) using a 2-45% in 45 min 0,1%TFA/ACN+0,1%TFA gradient. The identity of the peptide was confirmed using MALDI-MS (4800 MALDI TOF/TOF, Sciex)

Expression and purification of Scc1 fragment

A codon optimized gene for residues 252-420 of the human cohesin kleisin subunit (Uniprot O60216) was synthesized with a 3C protease site, an N-terminal cysteine and a C329S mutation, and cloned into the BamHI and EcoRI sites of pGEX-6P-1. The resulting plasmid pED145 was transformed into Escherichia coli ER2566 cells (New England Biolabs, fhuA2 lacZ::T7 gene1 [lon] ompT gal sulA11 R(mcr73::miniTn10--TetS)2 [dcm] R(zgb-210::Tn10--TetS) endA1 Δ(mcrCmrr)114::IS10) containing a Rosetta™-2 plasmid (Novagen), and selected on LB media containing 100 mg/ml ampicillin and 34 mg/ml chloramphenicol. For expression, overnight pre-cultures were diluted 1:100 into 1 liter prewarmed LB with fresh antibiotics, cultures were grown in baffled flasks shaking at 200 rpm at 37°C until the $OD_{600}$ reached ~0.6, and expression was induced by the addition of IPTG to a final concentration of 1 mM. After 3 hours of induction the cells were harvested by centrifugation (10 min, 4000 rpm, JLA8.1000 rotor), washed in 50 ml PBS, and resuspended in buffer A (50 mM Tris/HCl pH 7.5, 200 mM NaCl, 5% (w/v) glycerol, 0.05 mM TCEP). Cells were lysed using a French Press (Constant Systems) at 20 kpsi, 4°C and unbroken cells, debris and aggregates were pelleted in a Ti45 rotor (30 min, 40.000 rpm, 4°C). The lysate was applied to 2 ml prewashed glutathione resin (Thermo Fisher 25237) and incubated for one hour while rotating at 4°C. Subsequently, the resin was washed with 50 ml of buffer A and finally the kleisin fragment was eluted in 15 ml of buffer A supplemented with homemade 3C protease. The kleisin fragment was concentrated using a Vivaspin centrifugal concentrator (10 kDa cut-off), fluorescently labelled with Alexa Fluor™ 647 C2 Maleimide (Thermo Fisher A20347) and further purified by size exclusion chromatography (SEC) on a Superdex 200 Increase 10/300 column pre-equilibrated with buffer A.

Synthesis of dsDNA construct for fluorescence-based loop extrusion experiments

The synthesis of the dsDNA construct based on λ phage DNA was described previously (*4*). Briefly, single-stranded DNA ends of λ phage DNA (D1521, Promega) were filled with DNA polymerase I large fragment (M0210, New England Biolabs) lacking exonuclease activity, in the presence of dTTP, dCTP, dGTP, and biotin-14-dATP (19524016, Invitrogen). The DNA construct was purified from excess nucleotides and polymerase using a PCR clean-up kit (20021, Qiagen) and stored at –20°C until use. Once an aliquot was thawed, it was kept at 4°C.

Synthesis of dsDNA construct for magnetic tweezers loop extrusion experiments

Singly biotinylated, linear dsDNA constructs with a length of 1.5 kbp were synthesised via PCR and enzymatically ligated to digoxigenin-enriched DNA handles, as described previously (*4*). Briefly, a biotinylated DNA fragment of 1.5 kbp length was produced by using biotin-labeled forward and reverse primers that contain a BsaI restriction site on pBluescript II SK+. To create the digoxigenin-enriched handle, a 485 bp fragment from pBluescript II SK+ (Stratagene, Agilent Technologies Inc., USA) was amplified by PCR in the presence of 1:5 digoxigenin-11-dUTP:dTTP (Jena Bioscience, Germany). Prior to ligations of the DNA fragment and handle, the amplicons were digested with the non-palindromic restriction enzyme BsaI-HFv2 (New England Biolabs, UK). The ligation of the DIG-handle and biotinylated DNA fragment was carried out overnight using T4 DNA ligase (New England Biolabs, UK). The final dsDNA construct was cleaned up from the access of handle by running on a 1% agarose gel and extracting the dsDNA construct using a gel purification kit (A9282, Promega) and stored at -20°C until use.

ATPase assay

20 nM recombinant human cohesin tetramer, 50 nM recombinant human NIPBL-Mau2, and 10 ng/μl λ-DNA were incubated in ATPase reaction buffer (final composition: 25 mM $NaH_2PO_4/Na_2HPO_4$ pH 7.5, 50 mM NaCl, 2.5 mM $MgCl_2$, 1 mM DTT, 0.1 mg/ml BSA, 2 mM ATP, and 10 nM $^{[\gamma\text{-}32P]}$ATP (Hartmann Analytic; SRP-501) at 37°C. Reactions were stopped by adding 1% SDS and 10 mM EDTA in a 10 min interval with 40 min endpoint. Where indicated, reactions were supplemented with CTCF peptide (freshly dissolved in 50 mM $NaH_2PO_4/Na_2HPO_4$ pH 7.5, 100 mM NaCl, 5% glycerol). Reaction products were separated on polyethyleneimide plates (Sigma; 1055790001) by thin-layer-chromatography using 0.75 M $KH_2PO_4$ (pH 3.4), analysed by phosphor imaging with a Typhoon Scanner (GE Healthcare), and quantified using ImageJ. The measured ATPase rates were normalised to the ATPase rate of cohesin in absence of CTCF fragments of the respective experiment day.

Synthesis of 5(6)-Carboxyfluoresceine-labeled KTYQR peptide

Peptide was synthesised on a Liberty Blue peptide synthesiser (CEM) using standard Fmoc chemistry. For each amino acid cycle, 4min coupling with DIC/Oxyma was performed. N-terminal 5(6)-Carboxyfluoresceine was added under the same conditions as an amino acid. Peptide was purified on a Phenomenex Luna C18(2) using a 2-45% in 45 min 0,1%TFA/ACN+0,1%TFA gradient. The identity of the peptide was confirmed using MALDI-MS (4800 MALDI TOF/TOF, Sciex)

Single-molecule DNA loop extrusion assays

LE experiments and imaging were performed as described previously (*15*). LE reactions by human cohesin were carried out in a buffer containing 40 mM Tris-HCl pH 7.5, 30 mM NaCl, 2.5 mM $MgCl_2$, 2.5% D-glucose, 2 mM Trolox, 10 nM catalase, 18.75 nM glucose oxidase, 100 nM Sytox Orange, 0.5 mg/ml BSA, 1 mM DTT, 1 mM ATP with 100 pM cohesin and 200 pM NIPBL-Mau2 at 37°C. For experiments with NIPBL-ΔN-A550/JF646, 100 pM cohesin, 15 pM NIPBL-ΔN-A550, and 15 pM NIPBL-ΔN-JF646 was used with 25 nM Sytox Green instead of Sytox Orange. 2 µM peptide was added were indicated. All data were acquired using an exposure time of 200 ms for LE experiments without double-labeled NIPBL-ΔN and using an exposure time of 300 ms in experiments with double-labeled NIPBL-ΔN.

Quantification and segmentation of kymographs

Kymographs were quantified and segmented as described in (*15*). In brief, images were cropped and processed by a median filter with a window size of 5 frames and the background was subtracted using a Top-Hat filter of size 10 pixels. The intensity of the DNA loop is normalised to the intensity along the entire DNA molecule and multiplied by the known length of the DNA molecule (48.5 kbp). The DNA loop position was determined as the relative position of the loop from one end of the DNA. Segmentation of cohesin-mediated LE phases was performed semi-automatically under assistance of a change point detection algorithm that was subsequently manually curated since an entirely automatic segmentation was not robust and returns spurious results on some traces or on parts of traces. After filtering, the change point detection was performed using a window-based change point detection algorithm (*69*), which were subsequently merged and classified based on the error estimation of the assay (*15*, *26*). The direction of each LE and DNA loop slipping phase was subsequently annotated by subjecting the DNA loop position over time within each segment to a linear regression and classifying the segment as direction1/direction2 based on the slope of the linear fit. Note that the nomenclature of the direction is arbitrary. Here, the first direction to which the loop travels is called direction 1.

A LE trace was labeled as unidirectional if all LE phases in the trace pointed in the same direction and as bidirectional otherwise. The frequency of LE phases was computed by dividing the number of LE phases by the time between the start of the first and last LE phase of every trace. The direction switching frequency was computed by dividing the number of direction switches by the time between the start of the first and last LE phases of every trace.

The displayed DNA loop size and position traces (Fig. 1C) were filtered using a Savitzky–Golay filter with a window length of 51 frames (5.1 s) and order 1 (implemented as part of the *scipy* package (*70*)).

To analyse the relationship between NIPBL-ΔN exchange and cohesin's extrusion direction switching, we monitored instances where the fluorescence intensity of one label was replaced by another, or where the intensity of one label disappeared and reappeared at least two frames later. The intensity of NIPBL-ΔN-A550/JF646 was calculated as the average intensity in the respective channel at the loop position, using a 5-pixel window around the loop centroid. Background intensity was subtracted by averaging the intensity in a region adjacent to the cropped DNA molecule, free from non-specific fluorescently labeled molecules during the loop lifetime.

We determined the number of bleaching steps by plotting the fluorescence intensity at the loop over time and counting the stepwise decreases to the background level, assisted by a hidden Markov model (HMM) analysis (*71*). Only NIPBL-ΔN complexes associated with a DNA loop were considered. The NIPBL-ΔN residence time (Fig. S3C) was calculated as the interval between the disappearance and preceding appearance of a NIPBL-ΔN molecule.

To correlate NIPBL-ΔN exchanges with extrusion direction changes, we counted direction changes during periods when NIPBL-ΔN dissociated from cohesin. Conversely, we also calculated the fraction of events where a direction change occurred concomitantly with a NIPBL exchange (Fig. 2F-G).

Measuring DNA loop extrusion activity and steps with magnetic tweezers

The magnetic tweezers instrument and experimental methodology used in this study was previously described in detail (*47*, *49*). Bead x, y, z position tracking was achieved with a spatial resolution of ~2 nm (*47*). The concentration of cohesin was 100 pM together with 250 pM NIPBL-Mau2; the varying concentrations of the different CTCF fragments that were additionally added are denoted within the results description and Figures. Experiments were conducted at 22.3°C in a buffer containing 40 mM Tris pH 7.5, 40 mM NaCl, 2.5 mM $MgCl_2$, 1 mM DTT, 1 mM ATP, 0.05% Tween-20, 0.25 mg/ml BSA. Experiments with yeast condensin were conducted with 1.5-2 nM condensin at 22.3°C in a buffer containing 40 mM Tris pH 7.5, 40 mM NaCl, 2.5 mM $MgCl_2$, 1 mM DTT, 1 mM ATP, 0.25 mg/ml BSA. Data was acquired for 12 min after protein flush-in and lowering the applied force from 7 pN to 0.3 pN.

Datasets were processed using custom-written Igor v6.37-based scripts, removing traces that showed surface-adhered magnetic beads, ruptured tethers, as well as tethers containing more than one dsDNA (*47*, *49*). The bead Z-positions of all traces conducted at identical conditions were pooled and filtered to 1 Hz (moving average) for subsequent quantitative LE-step analysis.

Analysis of MT traces

The step-finding procedure is based on the quantitative method described by Smith in 1998 for detecting edges in noisy time-series data (*72*). The algorithm was first made accessible for utilisation in the Python programming environment within the framework of the stepfinder Python package developed for the identification of steps in one-dimensional data with low signal-to-noise ratio (*73*) and subsequently refined to suit the specific requirements of single-molecule force spectroscopy experiments.

In brief, the algorithm enables the identification of steps with consistent reliability despite variable noise levels in the data as follows: at each time point $\boldsymbol{x_i}$, the difference between the forward and backward estimation $\boldsymbol{\Delta_i = X_{i+} - X_{i-}}$ of a Chung-Kennedy filter (*74*) applied to the time-series data is compared to the local root-mean-square (RMS) noise, calculated from the weighted moving variances $\boldsymbol{w_{i\pm} s_{i\pm}}$ over the same window:

$$\boldsymbol{Y_i \frac{\Delta_i}{\sqrt{w_{i+}s_{i+} + w_{i-}s_{i-}}}},$$

where the weights $\boldsymbol{w}$ correspond to the normalised switching factors of the Chung-Kennedy filter and are defined as

$$\boldsymbol{w_{i\pm} = \frac{s_{i\pm}^{-p}}{s_{i\pm}^{-p} + s_{i-}^{-p}}},$$

with $\boldsymbol{p}$ being the nonlinearity factor (edginess) proposed by Chung and Kennedy. In practice, $\boldsymbol{p = 20}$ was chosen.

The noise component in the denominator is a function of the experimental conditions, e.g. magnetic tweezer force and DNA length. It consists of the weighted moving variance calculated to the mean of the data and averaged over a window and does not exhibit peaks at step locations, unlike the standard deviation. When an edge is encountered in the data, the output function $\boldsymbol{Y}$ responds with a triangular waveform peaking at the location of the edge itself.

The benefit of this edge detector is that its probability distribution is known, allowing for the calculation of confidence limits for the presence and absence of a step at each time point (*72*).

This relationship is leveraged to define an initial threshold for the detection of a peak in $\boldsymbol{Y}$

$$\boldsymbol{y_c} = \frac{\mathbf{2}}{\mathbf{3}} \cdot \frac{\text{minimum step size}}{\text{local noise}} \quad (\mathbf{1})$$

which ensures that the probability $\boldsymbol{P_T}$ for the detection of a true step (efficiency) is maintained at a constant level.

It can be shown that assuming an SNR of 1 and thus choosing $\boldsymbol{y_c = 2/3}$ leads to the detection of steps with an efficacy of 95%.

In addition, a downstream rejection procedure was applied to verify whether Eq. 1 is satisfied for each individual detected step. For this purpose, the local step noise was computed within an interval around the index of the event, which was determined by the minimum allowed step spacing corresponding to two times the filter window. Based on this noise component and $\boldsymbol{y_c}$, a dynamically adapting acceptance threshold value (in units of the input signal) for the current slice of the time trace was back-calculated corresponding to the minimum allowed step size in Eq. 1.

Above-threshold steps were considered valid, while below-threshold steps were rejected and the neighboring plateaus were merged. After each rejection, the quality of the remaining events was reassessed in an iterative procedure until the stepfinder output consisted of valid steps only (Fig. S1C,D). To address the problem that statistical predictors such as the probability of true and false outputs and location accuracy of the stepfinder are also a function of the filter window width $\boldsymbol{W}$, the detector was run with a sequence of 50 window widths between 0.5 s and 5.0 s and the results were analysed. After determining the value $\boldsymbol{W_{opt}}$ which yielded the minimum standard deviation of the step mass (step size over noise for each event) and the maximum ratio of valid over rejected steps, the final window size was conservatively set to $\frac{4}{5}\boldsymbol{W_{opt}}$ and the final step-finding analysis was conducted.

This procedure, named *LowForceAutoStepfinder* (LFAS) was evaluated against a previously employed method to identify steps in MT traces of SMC-mediated LE (*AutoStepfinder*) (*47*). To this end, we computed the true positive rate (TPR) of detected steps using step validation data as conducted previously (*47*). In brief, 1.5 kbp-long dsDNA was tethered between the glass surface and magnetic bead as described above and a force of 0.3 pN or 1 pN was applied. The piezo holding the objective was set to step up or down every 10 seconds for a range of step sizes (Fig. S1E,F). Both stepfinder algorithms were applied to these traces. These datasets were then compared to the known steps from the piezo motion. In line with the previous stepfinder evaluation (*47*), a step detection was judged correct if there was a piezo step nearby within 10% (~1 s) of the expected dwell time and 30% of the expected step size. We found that the LFAS has a higher TPR at smaller forces (in particular at 0.3 pN used in this study) and smaller induced step size (Fig. S1G) and was thus used throughout this study.

Fitting of activity reduction and reversal/consecutive step ratio data

To fit the activity reduction in dependence of the peptide concentrations from MT experiments, the correlation between the peptide concentrations and the activity reduction data was first assessed using Spearman's correlation coefficient (function *scipy.stats.spearmanr* of the *scipy* package (*70*)). If the correlation coefficient was below 0.8, the data points were not subject to a fit, but described by a horizontal line with a height equal to the average value of the data points and the error bar equals the ± SD of the data points (e.g. CTR and NTR+condensin data in Fig. 1M). If the Spearman's correlation coefficient exceeded 0.8, a piecewise linear model was fitted to the data that consisted of a horizontal line up to a concentration $\boldsymbol{x_0}$ and a line with a negative slope thereafter in logarithmic space (e.g. NTR data in Fig. 1I), i.e.

$$\begin{cases} \boldsymbol{B} \text{ if } \boldsymbol{x} \le \boldsymbol{x_0} \\ \boldsymbol{B} - \boldsymbol{m}\,\mathbf{log}(\boldsymbol{x}/\boldsymbol{x_0}) \text{ if } \boldsymbol{x} > \boldsymbol{x_0} \end{cases}$$

where $\boldsymbol{B}$ is an offset, m denotes the slope of the line in logarithmic space, and $\boldsymbol{x_0}$ denotes the concentrations at which the horizontal line transitions into the line with a negative slope.

To fit the ratio of reversal to consecutive steps in dependence of peptide concentrations, the same approach was used, but the data points were fitted to the following function if the Spearman's correlation coefficient exceeded 0.8 (e.g. YDF data in Fig. 2K):

$$\boldsymbol{y}(\boldsymbol{x}) = \boldsymbol{B} + \frac{\boldsymbol{L}}{\mathbf{1} + (\boldsymbol{x}/\boldsymbol{x_0})^{-\mathbf{1}/\boldsymbol{k}}}$$

This function is a logistic function with $\boldsymbol{x}$ and $\boldsymbol{x_0}$ on a logarithmic scale. $\boldsymbol{L} + \boldsymbol{B}$ denotes the supremum of the function, and $\boldsymbol{k}$ is the logistic growth rate.

To provide a simple and robust description of the observed increase in reversals and the decrease in consecutive steps as a function of peptide concentrations, an exponential model was applied if, again, the Spearman's correlation coefficient exceeded 0.8:

$$\boldsymbol{y}(\boldsymbol{x}) = \boldsymbol{A} \cdot \boldsymbol{exp}(-\boldsymbol{kx}) + \boldsymbol{B}$$

where the amplitude $\boldsymbol{A}$ indicates the magnitude of change, and $\boldsymbol{k}$ is the convergence rate describing how quickly the system approaches its asymptotic value $\boldsymbol{B}$, which represents either a maximum (for $\boldsymbol{A} < \mathbf{0}$) or a baseline (for $\boldsymbol{A} > \mathbf{0}$).

Fluorescence Correlation Spectroscopy (FCS)

Fluorophore diffusion measurements were conducted using version 1 coverslips on a Picoquant Microtime 200 microscope, operated with Symphotime software at room temperature. A 60x Olympus UPSLAPO 60XW water immersion objective (working distance: 280 µm, numerical aperture: 1.2) was used to focus 485 nm (used for experiments with Cy3-labeled dsDNA) and 530 nm lasers (used for experiments with 5(6)-Carboxyfluoresceine-labeled KTYQR peptide). Prior to the experiment, the molecular brightness of Atto488 or Atto550 fluorophore solutions was optimised by adjusting the objective's correction collar. Emission light was directed through a 50 µm pinhole, split by a dichroic mirror, and filtered using a 520/35 optical band pass filter (Semrock) and a 563/9 band pass filter (Chroma). Fluorescence emission was collected by single-photon avalanche-diode detectors (PD5CTC and PD1CTC, Micro Photon Devices, Bolzano).

To examine affinity changes of $STAG1^{Scc1}$ to DNA in dependence of the YDF peptide, 10 nM of a Cy3-labeled 40 bp dsDNA (3'-CGTAGTCGGAGATGCGATTTGCATACCACCAG CGTAGTCG-5'), 150 nM $Scc1^{251-420}$, 15 nM STAG1, and the indicated excess of YDF peptide over STAG1 were incubated in a buffer containing 10 mM HEPES pH 7.5, 25 mM NaCl and 2.5 mM $MgCl_2$ for 30 min at room temperature. $Scc1^{251-420}$ was omitted where indicated and an equal volume of $Scc1^{251-420}$ storage buffer was added instead. ADA peptide was used instead of YDF peptide where indicated. To examine binding of 5(6)-Carboxyfluoresceine-labeled KTYQR peptide to STAG1, increasing concentrations of STAG1 were incubated with 20 nM labeled KTYQR peptide in a buffer containing 10 mM HEPES pH 7.5, 25 mM NaCl and 2.5 mM $MgCl_2$ for 30 min at room temperature.

For fitting of the FCS curves (using PAM (*75*)), the size of the confocal volume was determined from measurements of the free dye by fitting a single-component diffusion model with triplet state (a diffusion constant of 297 $\mu m^2/s$ was used (*76*)). The axial and lateral sizes of the confocal volume were fixed for further analysis.

FCS curves of Cy3-labeled dsDNA binding to STAG1 were fitted with a single-component diffusion model with triplet state. The resulting diffusion constant was in good agreement with the theoretical diffusion constant of freely diffusing DNA (MW 26 kDa) and of dsDNA in complex with STAG1, $Scc1^{251-420}$, and YDF peptide (combined MW 182 kDa). Because the diffusion constant between the free and bound DNA differs only by a factor ~2, a two-component diffusion model with triplet state was overfitting the data.

FCS curves of 5(6)-Carboxyfluoresceine-labeled KTYQR peptide were fitted with a two-component diffusion model with triplet state with fixed diffusion constants of the free peptide (MW 2 kDa) and the KTYQR peptide in complex with STAG1 (combined MW 139 kDa). The relative amplitude of the bound and unbound fraction is shown in Fig. 3K.

### Conservation score annotation

Conservation scores were computed using the ConSurf webserver (consurf.tau.ac.il) (*77*, *78*).

### AlphaFold predictions and structure analyses

Structure predictions of cohesin subunits and CTCF fragments were performed with AlphaFold v2 (*59*, *60*), downloaded from *https://github.com/google-deepmind/alphafold*. We used the default parameters except that all models were relaxed with a maximum number of 20 outer and 200 inner iterations. The sequences of the individual cohesin and CTCF fragments are listed in Table S2 and the individual chains as input to the AlphaFold-multimer pipeline are listed in Table S3. After the structure prediction, individual predictions of a combination of cohesin subunits and CTCF fragments were aligned on the NIPBL, STAG1, STAG2, or Smc3, respectively, using custom scripts using *PyMOL* (*79*). The structures were also rendered in PyMOL. The structural similarity of individual amino acids within CTCF fragments #1-3 (and scrambles thereof) was computed as follows: the C$\boldsymbol{\alpha}$ coordinates were extracted from the PDB files and the distance of each amino acid in prediction pi to the corresponding amino acid in prediction $\boldsymbol{p_j}$ was computed, where $\boldsymbol{i} \neq \boldsymbol{j}$. Subsequently, the median distance of each amino acid in the prediction $\boldsymbol{p_i}$ to all other predictions was computed and displayed as one individual line (the median distance of each amino acid in prediction $\boldsymbol{p_i}$ to all other predictions) in Fig. S6C.

### 3D polymer simulations

3D polymer simulations were performed as described by Banigan et al. (*80*), with small modifications. A genomic region comprising 5600 monomers (1 monomer = 2 kb) was simulated containing 8 repeating elements of TADs spanning 400, 100, and 200 monomers respectively. These 8 elements were averaged to generate the contact maps shown in Fig. S4F-H. Loop extruders switched their extrusion side and extruders underwent diffusion and slipping phases according to the experimentally observed rates (*15*). When extruders encountered a barrier (CTCF), extruders stopped with the *stall probability*. If stalled, extruders stalled on both sides with the *probability to stall symmetrically*, i.e. extruders were not allowed to switch extrusion directions anymore and extrude away from CTCF. Additionally, extruders either could or could not be stalled by CTCF during diffusion and slipping phases. 1D simulations were equilibrated and 3D polymer simulations were subsequently performed as before (*80*). Simulations were run on nodes with access to a NVIDIA Tesla V100S using DelftBlue DHPC (*81*).

Simulations were run for the following combination of parameters, totaling 338 simulations:

- Stall probability = [0, 0.01, 0.05, 0.1, 0.5, 0.6, 0.65, 0.7, 0.75, 0.8, 0.85, 0.9, 0.95]
- Probability to stall symmetrically = [0, 0.01, 0.05, 0.1, 0.5, 0.6, 0.65, 0.7, 0.75, 0.8, 0.85, 0.9, 0.95]
- CTCF can stall during diffusion and slipping phases = [yes, no]

From the resulting polymer configurations, contact maps and the contact frequency as a function of genomic distance *s* (P(s)) were computed from 4000 conformations using the openmmlib package (https://github.com/mirnylab/openmm-polymer-legacy) with a contact radius of 5 monomers. The simulated P(s) curves were quantitatively compared to experimental P(s) curves from wild-type Hap1 cells as reported by Haarhuis et al. (*23*) and the goodness of fit between simulated and experimental P(s) curves was computed as the geometric standard deviation of the ratio of simulated to experimental P(s) as before (*82*). Each parameter combination was ranked by its goodness of fit either for a fixed combination of the stall probability and whether CTCF can stall cohesin during diffusion and slipping phases (Fig. S4B,C) or overall (Fig. S4D,E).

Quantification and statistical analyses

Statistical tests were performed using the *scipy* module (*70*) or using GraphPad Prism 9. Details of statistical test performed on the presented data are stated in the figure captions.

### Supplementary Text

#### CTCF stalls loop extrusion by cohesin from both sides

To explore to what extent stalling of cohesin, rather than direction reversal, can explain Hi-C maps, we performed 3D polymer simulations (Methods) (*80*, *82*), varying two key parameters: i) the probability of CTCF stalling cohesin and ii) the likelihood that this stalling also prevents extrusion away from the encountered CTCF) (Fig. S4A). We ranked parameter combinations by the geometric standard deviation between simulated contact frequencies as function of genomic distance *s* (P(s)) and experimental P(s) curves (*23*, *82*). Simulations in which the majority of cohesins were prevented from switching direction upon encountering CTCF outperformed those with one-sided stalling, resembling experimental Hi-C data more closely (Fig. S4B,C). For example, the highest-ranking simulation (CTCF stalls cohesin with 95% probability and symmetrically blocks extrusion 75% of the time; Fig. S4F) produced fewer secondary CTCF-CTCF loops than its counterpart assuming one-sided stalling (Fig. S4G), aligning well with experimental Hi-C maps (*23*) (Fig. S4H). We also explored whether cohesin can be stalled by CTCF during diffusion and slipping phases (and not only during active extrusion phases) but this had neglectable impact (Fig. S4D,E; Methods).

Together, these results support the conclusion that CTCF NTR largely inhibits cohesin upon encounter, rarely allowing extrusion away from CTCF. Instead, CTCF-CTCF loops may be formed *in vivo* by dimerisation of cohesin$^{STAG2}$ (*83*).

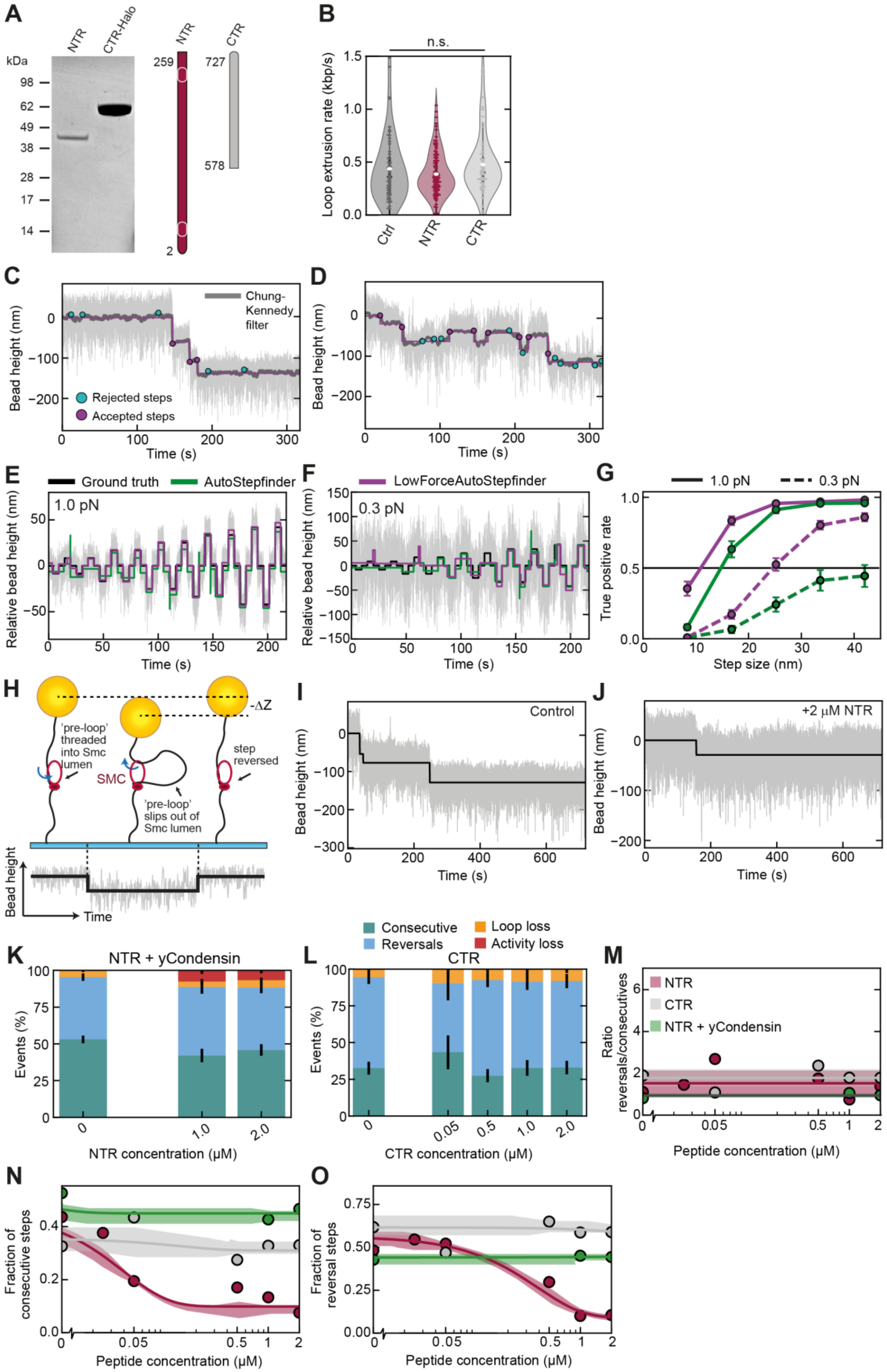
A
NTR
CTR-Halo
kDa
98
62
49
38
28
17
14
NTR
CTR
259
727
578
2
B
n.s.
Loop extrusion rate (kbp/s)
Ctrl
NTR
CTR
C
Chung-Kennedy filter
Bead height (nm)
Rejected steps
Accepted steps
Time (s)
D
Bead height (nm)
Time (s)
E
Ground truth
AutoStepfinder
1.0 pN
Relative bead height (nm)
Time (s)
F
LowForceAutoStepfinder
0.3 pN
Relative bead height (nm)
Time (s)
G
1.0 pN
0.3 pN
True positive rate
Step size (nm)
H
-ΔZ
'pre-loop' threaded into Smc lumen
SMC
'pre-loop' slips out of Smc lumen
step reversed
Bead height
Time
I
Control
Bead height (nm)
Time (s)
J
+2 µM NTR
Bead height (nm)
Time (s)
Consecutive
Reversals
Loop loss
Activity loss
K
NTR + yCondensin
Events (%)
NTR concentration (µM)
L
CTR
Events (%)
CTR concentration (µM)
M
NTR
CTR
NTR + yCondensin
Ratio reversals/consecutives
Peptide concentration (µM)
N
Fraction of consecutive steps
Peptide concentration (µM)
O
Fraction of reversal steps
Peptide concentration (µM)

**Figure S1. SDS-PAGE of CTCF NTR and CTR, cohesin LE rate, development of *LowForceAutoStepfinder*, additional MT data. Related to Fig. 1**.
(**A**) Coomassie staining of CTCF NTR and CTR fragments after SDS-PAGE.
(**B**) LE rate of cohesin in presence of 2 μM of the indicated peptide. Statistical significance was assessed by a Kruskal-Wallis test. N = 57, 110, 61 from left to right.
(**C,D**) Sep detection output (magenta) for two representative MT trajectories at 0.3 pN.
(**E,F**) Example MT traces of piezo-induced $\Delta Z$ steps measured on 1.5 kbp DNA at 1.0 pN and 0.3 pN stretching force, respectively, serving as a means to evaluate the *LowForceAutoStepfinder* (LFAS, magenta) with the AutoStepfinder (AS (*47*), green).
(**G**) Evaluation of the step detection sensitivity and detection limits of the step-finding methods on piezo-induced stepping trajectories using the true positive rate (Methods).
(**H**) Illustration of a failed attempt of an SMC protein complex to conclude a LE step cycle with an exemplary MT trace (bottom) displaying a reversal step in our working model. DNA is reeled through the Smc lumen which causes a step. Failure to stabilise the pre-loop and to merge it with the already-extruded loop causes reversal of the bead height to the original level.
(**I,J**) Example MT trace of cohesin in the absence (I) and presence (J) of 2 μM CTCF NTR. The light grey line show the raw data and the solid black line is the resulting stepfinder fit.
(**K**) Fractions of stepping events classified as consecutive LE steps, reversals, and loop loss for MT traces of DNA loop extrusion by yeast condensin with increasing concentrations of CTCF NTR. Error bars denote 95% binomial confidence intervals (CI). N = 1581, 454, 636 steps from 171, 50, 77 tethers, respectively, for increasing CTCF NTR concentration
(**L**) Same as (K) for LE by cohesin with in increasing concentrations of CTCF CTR. N = 445, 83, 466, 409, 609 from 114, 51, 62, 48, 96 tethers, respectively.
(**M-O**) The ratio of reversal and consecutive steps (M), fraction of consecutive (N), and fraction of reversal steps (O) with increasing concentrations of the indicated peptides from data as in Fig. 1I.

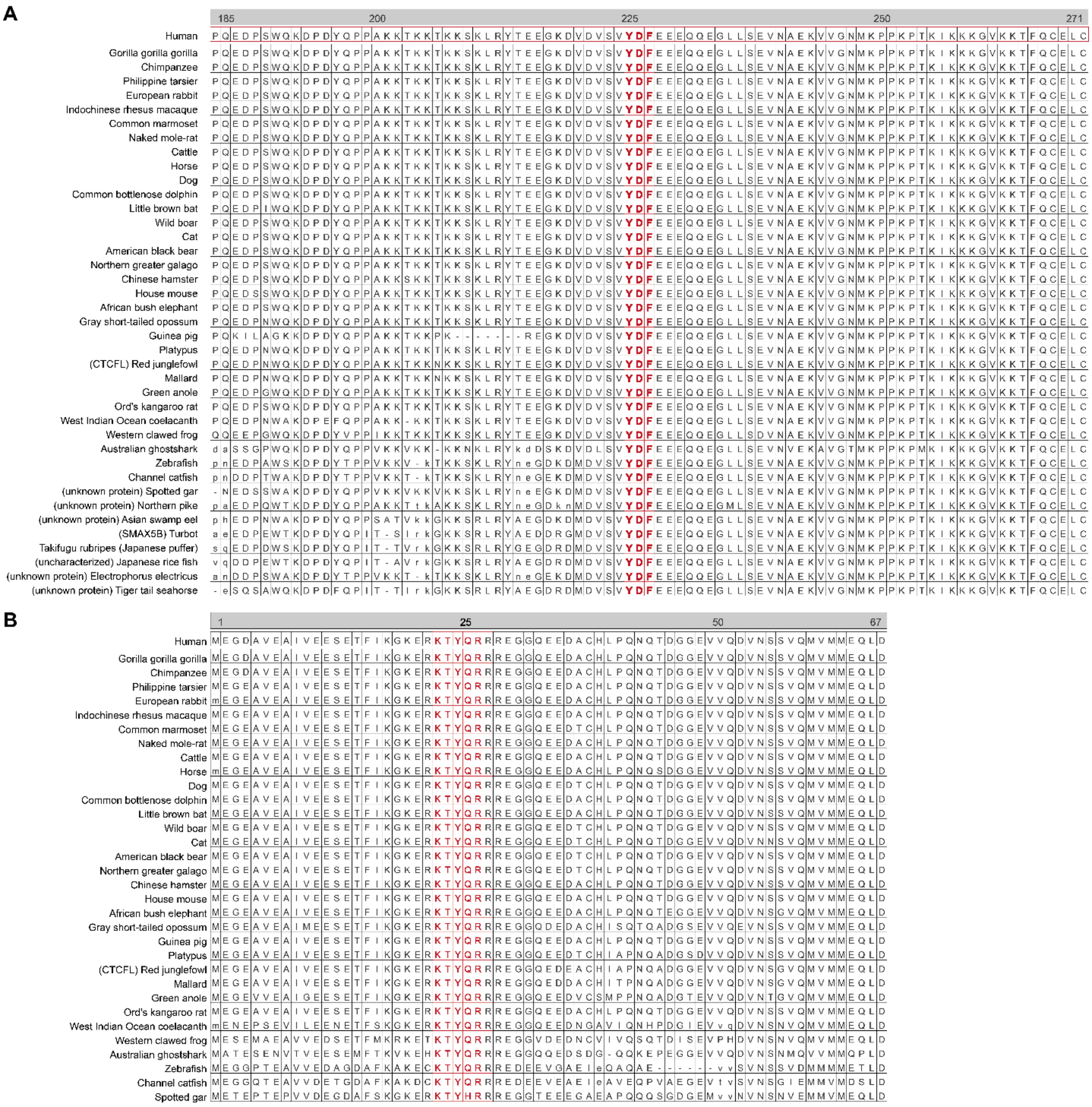

**Figure S2. Sequence homology assessment via SLiMSearch of CTCF fragments surrounding the YDF and KTYQR motifs of human CTCF. Related to Fig. 2 and 3.**

(**A**) SLiMSearch4 (*84*) of the YDF motif (motif regular expression used was [PFCAVIYL][FY][GDEN]F.{0,1}[DANE].{0,1}[DE] as described in (*33*)) with default parameters. Sequences were subsequentially aligned using ProViz (*85*) within the metazoa database.

(**B**) Same as (A) but surrounding the KTYQR motif (motif regular expression used was KTYQR).

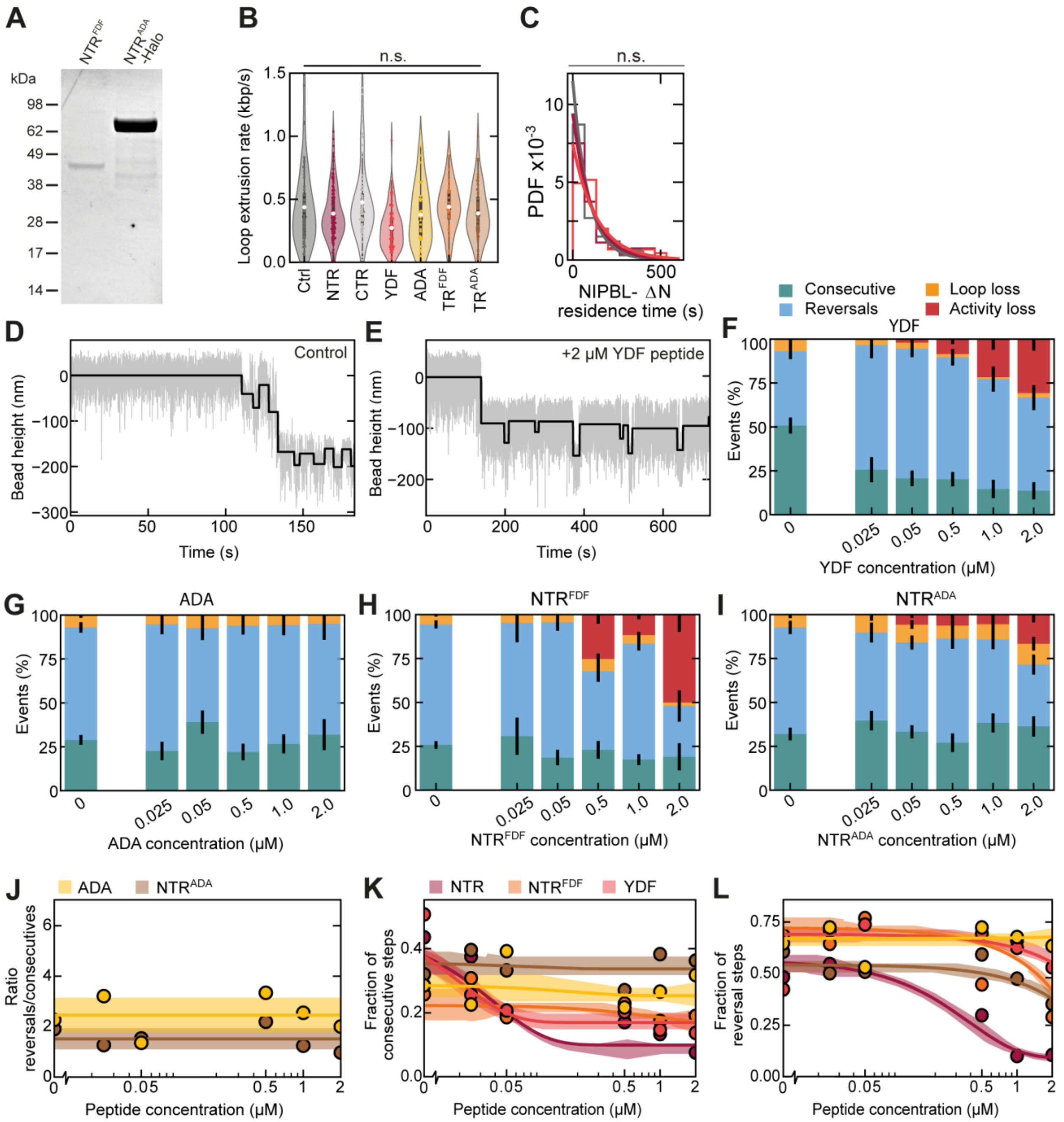


**Figure S3. SDS-PAGE of NTR$^{FDF}$ and NTR$^{ADA}$ and cohesin stepping activity in presence of the ADA peptide, NTR$^{FDF}$, and NTR$^{ADA}$.**

(**A**) Coomassie staining of CTCF NTR$^{FDF}$ and NTR$^{ADA}$ fragments after SDS-PAGE.

(**B**) LE rate of cohesin in presence of 2 μM of the indicated peptide. Statistical significance was assessed by a Kruskal-Wallis test. N = 57, 110, 61, 72, 32, 25, 57 from left to right.

(**C**) Distribution of NIPBL-ΔN residence times (N = 50, 67, 40 for control, NTR, and YDF, respectively). Statistical significance was assessed using a one-way ANOVA (n.s.: $p > 0.05$).

(**D-E**) Example MT trace of cohesin in the absence (D) and presence (E) of 2 μM YDF peptide. The light grey line denotes the raw data and the solid black line is the resulting stepfinder fit.

(**F**) Fraction of events classified as consecutive steps, reversals, and loop loss for MT traces of DNA loop extrusion by cohesin with increasing concentrations of YDF peptide. Bar heights denote the fraction of beads in the respective category and error bars denote 95% binomial confidence intervals (CI). N = 463, 203, 337, 344, 139, 132 steps from 155, 97, 133, 125, 41, 44 tethers, respectively, for increasing peptide concentration.

(**G**) Same as (F) with increasing concentrations of ADA peptide. N = 1060, 297, 237, 366, 357, 145 steps from 276, 79, 66, 72, 41, 77 tethers, respectively, for increasing peptide concentration.
(**H**) Same as (F) but with increasing concentrations of CTCF NTR$^{FDF}$. N = 1604, 104, 360, 202, 932, 50 steps from 432, 48, 69, 62, 223, 79 tethers, respectively, for increasing peptide concentration.
(**I**) Same as (F) but with increasing concentrations of CTCF NTR$^{ADA}$. N = 644, 333, 573, 253, 296, 225 steps from 142, 69, 134, 62, 72, 62 tethers, respectively, for increasing peptide concentration.
(**J-L**) The ratio of reversal and consecutive steps (J), fraction of consecutive (K), and fraction of reversal steps (L) with increasing concentrations of the indicated peptides from data as in Fig. 2I.

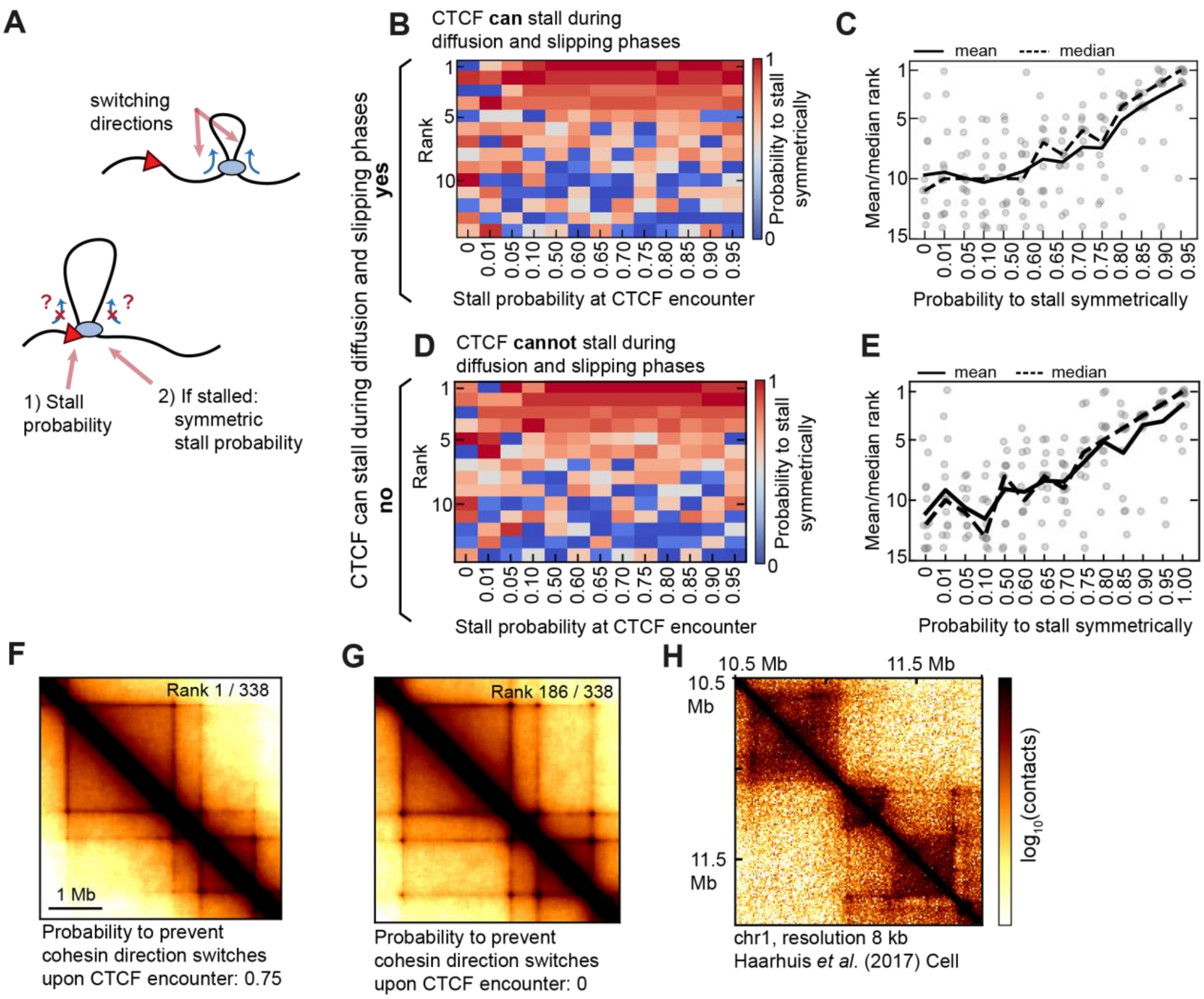


**Figure S4. Polymer simulations in which CTCF prevents cohesin to switch direction upon encounter recapitulate experimental Hi-C matrices**.

(**A**) Simulation setup: The probability to stall at CTCF upon encounter. If cohesin is stalled, cohesin stalls symmetrically, i.e. cannot switch directions after encountering a CTCF molecule, with a probability that is varied across simulation runs.

(**B**) Rank of individual simulations by their goodness of fit between simulated and experimental P(s) versus cohesin's stall probability at CTCF encounter. The heatmap is colored according to the simulation's probability to prevent cohesin switching directions upon being stalled on one side by CTCF. Cohesin can be stalled during diffusion and slipping phases.

(**C**) Mean and median rank of simulations versus probability to stall cohesin on both sides upon CTCF encounter. Individual data points correspond to different values of the stall probability. Cohesin can be stalled during diffusion and slipping phases.

(**D-E**) As for B,C but CTCF cannot stall cohesin during diffusion and slipping phases phases.

(**F**) Hi-C map of best-fitting simulation to the experimentally observed P(s) (probability to prevent cohesin direction switches upon CTCF encounter 0.75; block probability 0.95; diffusion and slipping can stall).

(**G**) Same as in (H) except that the probability to prevent cohesin direction switches upon CTCF encounter was set to 0.

(**H**) Experimental Hi-C matrix from wildtype Hap1 cells from Haarhuis *et al.* (*23*), extracted from HiGlass (*86*).

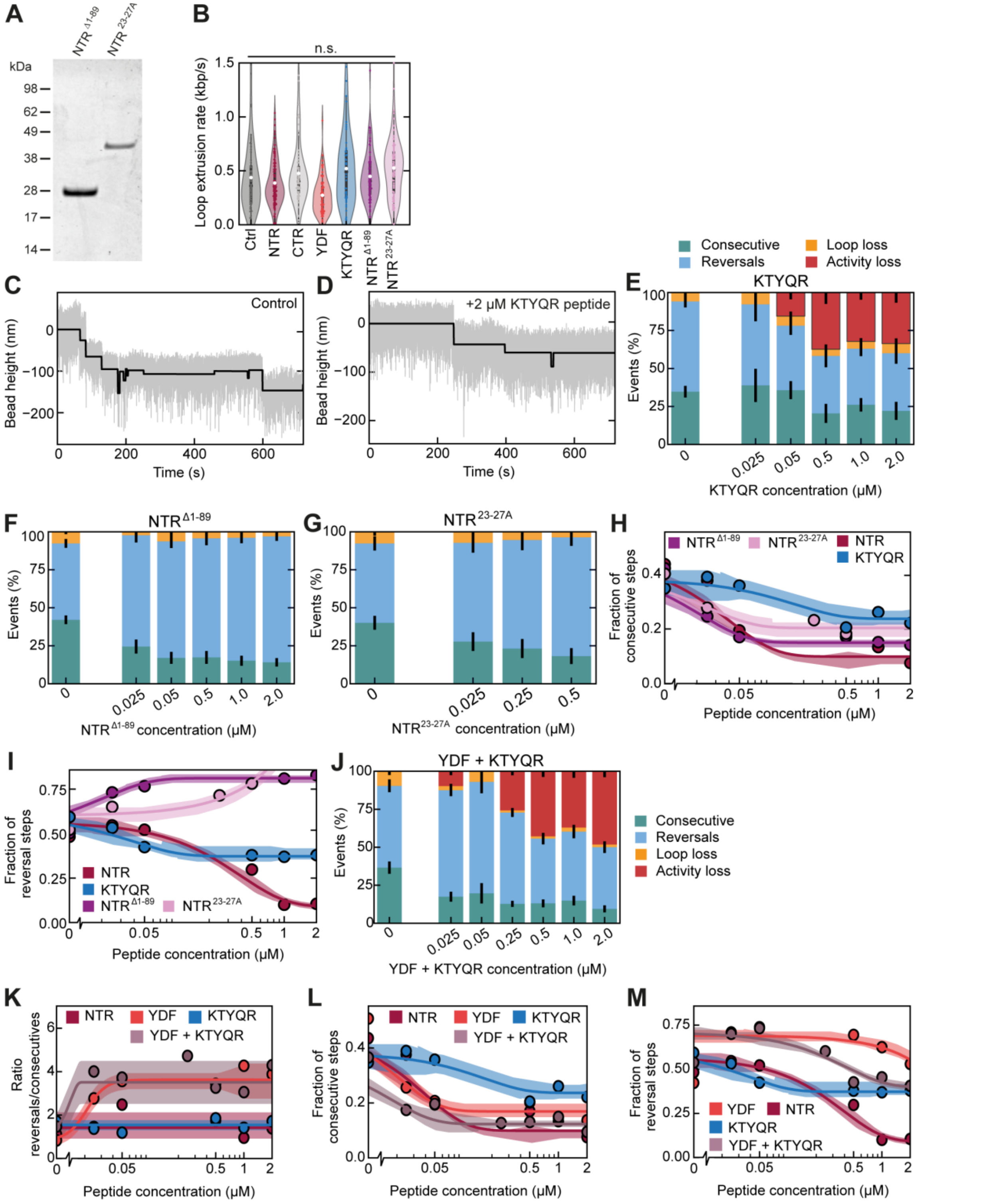


**Figure S5. SDS-PAGE of NTR$^{D1-89}$ and NTR$^{23-27A}$, cohesin stepping activity in presence of the NTR$^{D1-89}$ and NTR$^{23-27A}$ fragments.**

(**A**) Coomassie staining of CTCF NTR$^{D1-89}$ and NTR$^{23-27A}$ fragments after SDS-PAGE.

(**B**) LE rate of cohesin in presence of 2 µM of the indicated peptide. Statistical significance was assessed by a Kruskal-Wallis test. N = 57, 110, 61, 72, 46, 76, 79 from left to right.

(**C-D**) Example MT trace of cohesin in the absence (D) and presence (E) of 2 µM KTYQR peptide. The light grey line denotes the raw data and the solid black line is the resulting stepfinder fit.

(**E**) Fraction of events classified as consecutive steps, reversals, and loop loss for MT traces of DNA loop extrusion by cohesin with increasing concentrations of the KTYQR peptide. Bar heights denote the fraction of beads in the respective category and error bars denote 95% binomial confidence intervals (CI). N = 599, 90, 215, 101, 257, 126 steps from 295, 68, 90, 49, 185, 64 tethers, respectively, for increasing peptide concentration.
(**F**) As for (E) but with increasing concentrations of CTCF NTR$^{\Delta 1\text{-}89}$. N = 1155, 355, 429, 608, 911, 860 steps from 277, 70, 52, 94, 137, 154 tethers tethers, respectively, for increasing peptide concentration.
(**G**) Same as (E) but with increasing concentrations of CTCF NTR$^{23\text{-}27A}$. N = 444, 292, 522, 329 steps from 94, 37, 40, 38 tethers, respectively, for increasing peptide concentration.
(**H-I**) Fraction of consecutive (H), and reversal steps (I) with increasing concentrations of the indicated peptides from data as in Fig. 3F.
(**J**) Same as (E) but with increasing concentrations of YDF and KTYQR peptides. N = 544, 527, 300, 811, 381, 309, 329 steps from 242, 238, 81, 455, 307, 223, 315 tethers, respectively, for increasing peptide concentration.
(**L-M**) Fraction of consecutive (C) and reversal (D) steps with increasing concentrations of the indicated peptides from data as in Fig. 3J.
(**K**) Ratio of reversal and consecutive steps with increasing concentrations of the indicated peptides from data as in Fig. 3J.

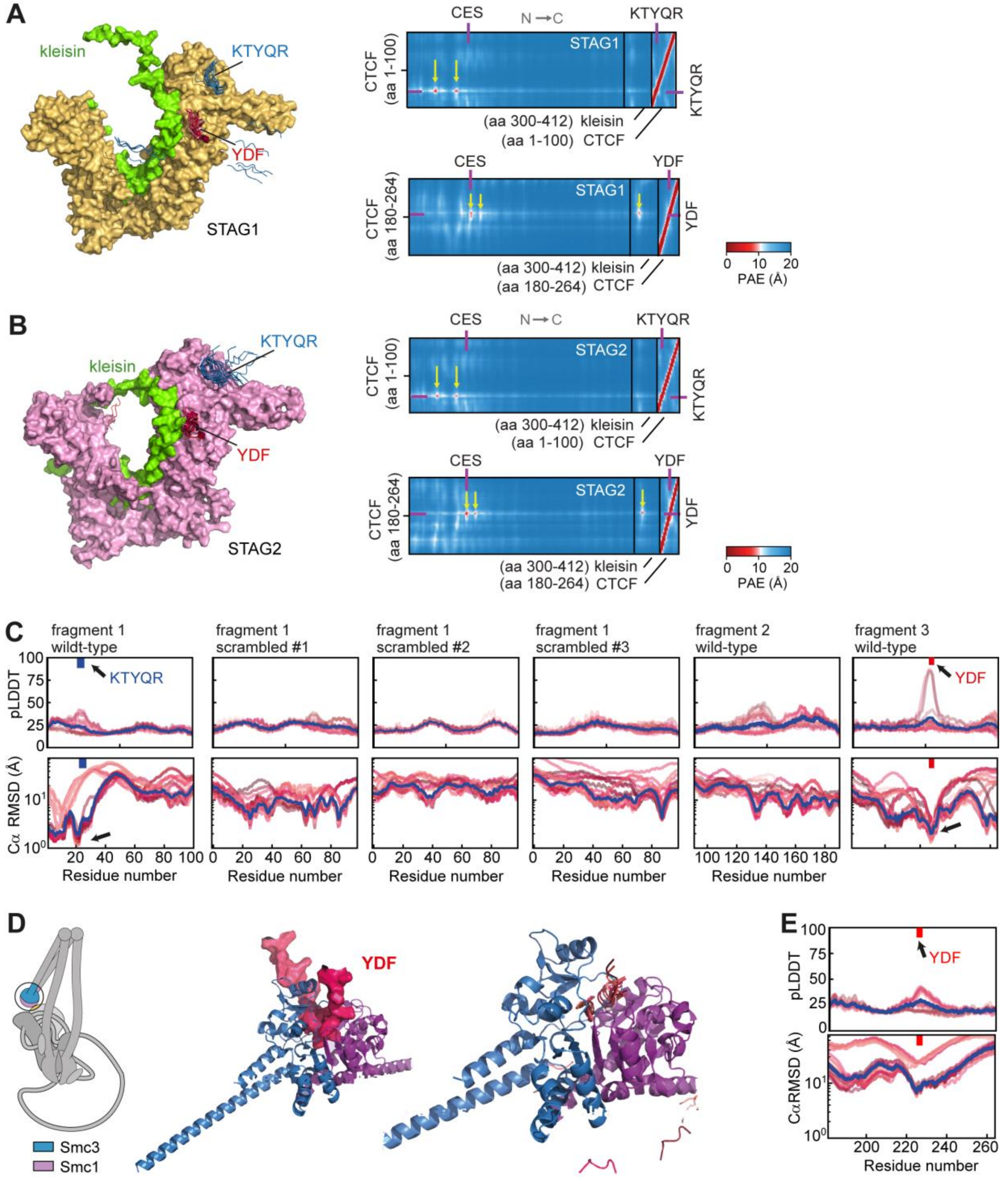


**Figure S6. AlphaFold predictions of cohesin subunits and fragments of CTCF**.

(**A**) AlphaFold predictions of STAG1 (left) with a kleisin fragment and a CTCF fragment spanning residues aa 1-100 containing the KTYQR motif and a CTCF fragment spanning residues aa 180-264 containing the YDF motif. The 25 predictions were aligned using PyMOL (*79*) onto the STAG1 structure prediction that were virtually identical to one another. Only the KTYQR motif (blue) or YDF motif (red) ± 4 aa is shown for each CTCF fragment chain for clarity. The placement of the KTYQR and YDF motif resulting from the 25 individual runs are shown individually. In 16/25 predictions, the KTYQR motif is placed N-terminally of the 'conserved essential surface' (CES (*33*, *38*, *87*)). In 25/25 predictions, the YDF motif is placed at the CES as observed previously. Right: Predicted Alignment Error (PAE)

maps of the three subcomplexes with low PAE scores between the YDF and KTYQR peptides (purple ticks) at the CES and a region located N-terminally of STAG1's CES on STAG1, respectively.
(**B**) Same as A but for STAG2 and maps for the YDF and KTYQR motifs.
(**C**) Predicted local distance difference test (pLDDT; scale from 0 to 100) values for 25 individual runs (red-shaded lines; blue line denotes the median) over CTCF fragment #1 (aa 1-100), 3 randomly scrambled versions of this sequence, CTCF fragment #2 (aa 90-190), and CTCF fragment #3 (aa 180-264). The positions of the KTYQR and YDF motif are annotated (blue and red arrows, respectively). Note that a low pLDDT value is expected since the CTCF NTR is predominantly intrinsically disordered (*88*). The bottom row shows the average distance of each amino acid within each CTCF fragment to the same amino acid in the same fragments of the other 24 runs. Values approaching 1 Å are only observed for the KTYQR and the YDF motif, indicating that independently placed motifs are (to some extent for the KTYQR motif) consistently placed at a single predicted binding site of STAG1 (and kleisin for the YDF motif).
(**D**) AlphaFold predictions of the Smc hinge and CTCF fragment #3 (containing the YDF motif). The 25 predictions were aligned on Smc3 and the 25 individual placements of the YDF motif are shown as red ribbons.
(**E**) As in (C) for CTCF fragment #3 in association with the Smc hinge.

**Table S1. Peptide sequences of CTCF fragments.** YDF and KTYQR motifs, as well as mutations thereof, are marked in bold.

| Peptide | Sequence | Position | Provider |
|---|---|---|---|
| **YDF** | DVSV**YDF**EEE | 222-231 | In-house |
| **ADA** | DVSV**ADA**EEE | 222-231 | In-house |
| **KTYQR** | FIKGKER**KTYQR**RREG | 17-32 | GenScript |
| **NTR** | EGDAVEAIVEESETFIKGKER**KTYQR**RREGGQEEDACHLPQNQT DGGEVVQDVNSSVQMVMMEQLDPTLLQMKT EVMEGTVAPEAEAAVDDTQIITLQVVNMEEQPINIGEL QLVQVPVPVTVPVATTSVEELQGAYENEVSKEGLAESE PMICHTLPLPEGFQVVKVGANGEVETLEQGELPPQEDP SWQKDPDYQPPAKKTKKTKKSKLRYTEEGKDVDVSV**YDF**EEE QQEGLLSEVNAEKVVGNMKPPKPTKIKK | 2-259 | In-house |
| **CTR** | AGPDGVEGENGGETKKSKRGRKRKMRSKKEDSSDSEN AEPDLDDNEDEEEPAVEIEPEPEPQPVTPAPPPAKKR RGRPPGRTNQPKQNQPTAIIQVEDQNTGAIENIIVEV KKEPDAEPAEGEEEEAQPAATDAPNGDLTPEMILSMMDR | 578-727 | In-house |
| **NTR$^{ADA}$** | EGDAVEAIVEESETFIKGKER**KTYQR**RREGGQEEDACH LPQNQTDGGEVVQDVNSSVQMVMMEQLDPTLLQMKT EVMEGTVAPEAEAAVDDTQIITLQVVNMEEQPINIGEL QLVQVPVPVTVPVATTSVEELQGAYENEVSKEGLAESE PMICHTLPLPEGFQVVKVGANGEVETLEQGELPPQEDP SWQKDPDYQPPAKKTKKTKKSKLRYTEEGKDVDVSV **ADA**EEEQQEGLLSEVNAEKVVGNMKPPKPTKIKK | 2-259 | In-house |
| **NTR$^{FDF}$** | EGDAVEAIVEESETFIKGKER**KTYQR**RREGGQEEDACH LPQNQTDGGEVVQDVNSSVQMVMMEQLDPTLLQMKT EVMEGTVAPEAEAAVDDTQIITLQVVNMEEQPINIGEL QLVQVPVPVTVPVATTSVEELQGAYENEVSKEGLAESE PMICHTLPLPEGFQVVKVGANGEVETLEQGELPPQEDP SWQKDPDYQPPAKKTKKTKKSKLRYTEEGKDVDVSV **FDF**EEEQQEGLLSEVNAEKVVGNMKPPKPTKIKK | 2-259 | In-house |
| **NTR$^{\Delta 1\text{-}89}$** | VDDTQIITLQVVNMEEQPINIGELQLVQVPVPVTVPV ATTSVEELQGAYENEVSKEGLAESEPMICHTLPLPEG FQVVKVGANGEVETLEQGELPPQEDPSWQKDPDYQPP AKKTKKTKKSKLRYTEEGKDVDVSV**YDF**EEEQQEGLL SEVNAEKVVGNMKPPKPTKIKK | 90-259 | In-house |
| **NTR$^{23\text{-}27A}$** | EGDAVEAIVEESETFIKGKER**AAAAA**RREGGQEEDACH LPQNQTDGGEVVQDVNSSVQMVMMEQLDPTLLQMKT EVMEGTVAPEAEAAVDDTQIITLQVVNMEEQPINIGEL QLVQVPVPVTVPVATTSVEELQGAYENEVSKEGLAESE PMICHTLPLPEGFQVVKVGANGEVETLEQGELPPQEDP SWQKDPDYQPPAKKTKKTKKSKLRYTEEGKDVDVSV **YDF**EEEQQEGLLSEVNAEKVVGNMKPPKPTKIKK | 2-259 | In-house |

**Table S2. Sequences of CTCF fragments and cohesin subunits used for AlphaFold predictions.** All subunits refer to the human sequence. The Smc1/3 head was assembled by the N- and C-termini of Smc1 and Smc3, respectively, linked by a flexible linker consisting of 32 Glycines (32xG).

| Name | Position |
|---|---|
| Scc1 fragment binding to NIPBL | 143-181 |
| Scc1 fragment binding to STAG1/2 | 301-413 |
| Scc1 fragment binding to Smc3 | 1-100 |
| Scc1 fragment binding to Smc1 | 531-631 |
| NIPBL | 1191-2641 |
| Mau2 | 1-613 |
| STAG1 | 86-971 |
| STAG2 | 77-1064 |
| Smc1 head | (1-240)-32xG-(994-1233) |
| Smc3 head | (1-201)-32xG-(978-1217) |
| Smc1 hinge | 481-711 |
| Smc3 hinge | 451-691 |
| CTCF fragment #1 | 1-100 |
| CTCF fragment #2 | 90-190 |
| CTCF fragment #3 | 180-264 |

**Table S3. Components of cohesin subunits used in AlphaFold predictions**

| Name | Components (nomenclature as in Table S2) |
|---|---|
| NIPBL-Mau2 | NIPBL, Mau2, Scc1 fragment binding to NIPBL |
| STAG1 | STAG1, Scc1 fragment binding to STAG1/2 |
| STAG2 | STAG2, Scc1 fragment binding to STAG1/2 |
| ATPase heads | Smc1 head, Smc3 head, Scc1 fragment binding to Smc1, Scc1 fragment binding to Smc3 |
| Hinge | Smc1 hinge, Smc3 hinge |
